# Dividing out quantification uncertainty enables assessment of differential transcript usage with limma and edgeR

**DOI:** 10.1101/2025.04.07.647659

**Authors:** Pedro L. Baldoni, Lizhong Chen, Mengbo Li, Yunshun Chen, Gordon K. Smyth

## Abstract

Differential transcript usage (DTU) refers to changes in the relative abundance of transcript isoforms of the same gene between experimental conditions, even when the total expression of the gene doesn’t change. DTU analysis requires the quantification of individual isoforms from RNA-seq data, which has a high level of uncertainty due to transcript overlap and read-to-transcript ambiguity (RTA). Popular DTU analysis methods do not directly account for the RTA overdispersion within their statistical frameworks, leading to reduced statistical power or poor error rate control, particularly in scenarios with small sample sizes. This article presents limma and edgeR analysis pipelines that account for RTA during DTU assessment. Leveraging recent advancements in the limma and edgeR Bioconductor packages, we propose DTU analysis pipelines optimized for small and large datasets with a unified interface via the diffSplice function. The pipelines make use of divided counts to remove RTA-induced dispersion from transcript isoform counts and account for the sparsity in transcript-level counts. Simulations and analysis of real data from mouse mammary epithelial cells demonstrate that the diffSplice pipelines provide greater power, improved efficiency, and improved FDR control compared to existing specialized DTU methods.

## Introduction

RNA sequencing (RNA-seq) has revolutionized biomedical research by enabling comprehensive profiling of the transcriptome, providing insights into gene expression regulation across diverse biological contexts, including cancer, immunology, and developmental biology. A common task in RNA-seq data analysis is to identify genomic features that have altered expression levels across conditions, such as treatments, disease status, or genotypes. Differential expression (DE) analysis has traditionally focused on genes as the primary units of expression [1]. However, genes often express multiple transcript isoforms (transcripts) via alternative splicing, a process in which gene exons are joined in different combinations, resulting in distinct messenger RNA products [2, 3, 4]. Recent computational and statistical developments now allow fast and accurate detection of differential transcript expression (DTE) [5, 6]. Yet, transcriptional changes resulting from alternative splicing rarely occur in isolation, as biological processes often affect multiple expressed transcripts of a gene simultaneously. Examples of such processes include alternative splicing via transcription start site variation and isoform switching via exon skipping [7]. These phenomena often occur in the context of cancer, where an oncogene transcript replaces a major transcript due to DNA damage or epigenomic modifications [8, 9]. It is therefore of key interest for biomedical researchers to identify those genes for which any differential splicing event has occurred, resulting in changes in the relative abundance of expressed transcripts for that gene between conditions.

Differential splicing can be assessed either at the level of exons via differential exon usage (DEU) or at the level of transcripts (RNA isoforms) via differential transcript usage (DTU). In DEU analyses, RNA-seq reads are aligned to a reference genome with a splice-aware aligner, reads are counted for exons, and statistical tests are conducted for changes in relative exon expression between experimental conditions [10]. Alternatively, reads might be counted for exon-exon junctions or for disjoint exon bins [11, 12]. In DTU analyses, RNA-seq reads are aligned to a reference transcriptome, transcript counts are estimated by quantification tools such as *Salmon* [13], *kallisto* [14], and *RSEM* [2], and tests are conducted for changes in the relative expression of the transcripts for each gene [15, 16]. The key step in transcriptome alignment is to determine which set of transcripts each sequence read is compatible with, where each possible set of compatible transcripts is called an *equivalence class* [14, 13]. A third way to assess differential splicing is to apply DEU statistical methods to the equivalence class counts instead of to exon counts [17]. Finally, it is also possible to test for DTU directly from the equivalence class counts instead of from transcript counts, although this approach requires complex Bayesian statistical models and long computation times [18]. All of these approaches have their challenges. DEU has the complication that exons do not have consistent start and end sites between transcripts and therefore are not consistently defined. Another issue is that sequence reads may be multiply assigned to more than one exon bin, causing inter-dependence between the exon counts. DEU applied to equivalence class counts avoids the problem of double-counting, but has to deal with a very large number of equivalence classes, and equivalence classes containing transcripts from more than one gene have to be ignored [17]. While all these approaches can test for differential splicing at the gene-level, only DTU can reveal which transcripts are being differentially used as a response to a given biological condition. This paper therefore focuses on DTU using estimated transcript counts.

The key statistical challenge that complicates the analysis of transcript counts is that they are estimated quantities and are highly overdispersed compared to exon-level or gene-level counts. The overdispersion arises from the fact that each sequence read is typically compatible with more than one reference transcript, a phenomenon that we call read-to-transcript ambiguity (RTA) [5]. Reads have to be assigned to transcripts probabilistically [2, 14, 13], and the assignment process tends to reinforce transcripts that are already highly expressed, leading to inflated variances for the estimated counts. The amount of overdispersion depends on the transcript annotation topology, and is greater for transcripts that are highly overlapping. The overdispersion arising from RTA not only increases the variability of the data but also, crucially, interferes with the global mean-variance relationships that allow gene-level RNA-seq analysis methods such as *edgeR* [19], *limma-voom* [20], and *DESeq2* [21] to be so powerful and accurate. We recently showed that the RTA overdispersions can be very accurately estimated from technical resamples generated by the transcript quantification tools [5, 6]. We showed that RTA induces quasi-Poisson overdispersion into the intra-sample transcript counts. The RTA overdispersions were shown to add technical variation to the mean-variance relationship of the transcript counts without affecting the biological coefficient of variation (BCV) [5, 6]. The fact that the RTA overdispersions are purely technical and also accurately estimated allows them to be, literally, divided out of the transcript counts. This process leads to divided counts that show the same negative binomial-type variation and global mean-variance relationships usually associated with gene-level analyses [5]. The divided-counts method was shown to dramatically improve the accuracy and efficiency of DTE analyses, in both large and small scale datasets, but has not yet been benchmarked for DTU.

*DEXSeq* [10], *DRIMSeq* [15], and *satuRn* [16] are popular software tools used to test for DTU. *DEXSeq* was originally developed to test for DEU but has been repurposed for DTU using transcript counts [12, 22]. *DEXSeq* starts by fitting negative binomial generalized linear models to the counts for each transcript, following the *DESeq2* method, in order to estimate the negative binomial dispersion parameter for each transcript. It then assesses differential usage by testing for interaction terms in gene-level generalized linear models. The gene-level linear predictor includes main effects for sample and transcript, so the interaction test may be considered as a generalization of the traditional G-test for independence in contingency tables [23, 24]. A test is conducted for each transcript, looking for differential usage of that transcript vs all other transcripts for the same gene. *DRIMSeq* assumes a Dirichlet-multinomial model for the transcript counts, treating the total gene count for each sample as fixed. The Dirichlet precision parameter is estimated for each gene and likelihood ratio tests are conducted for differential transcript count proportions. *satuRn* assumes a quasibinomial generalized linear model for the counts of each transcript, as a proportion of the total counts for that gene in each sample. Empirical Bayes moderation is applied to the quasi-binomial dispersion parameters, using *limma*’s *squeezeVar* function [25, 26]. Transcript-level t-statistics are computed for differential usage, and Efron’s local FDR approach [27] is used to obtain empirical p-values, which are then corrected for multiple testing.

None of the above DTU methods allow for technical overdispersion arising from RTA. They all estimate overdispersion only in terms of sample to sample variation arising from biological differences in transcript expression or transcript proportions. *DEXSeq*’s negative binomial model, and the multinomial and binomial models of *DRIMSeq* and *satuRn*, all assume that the transcript counts will be Poisson distributed under repeat sampling for an individual sample, whereas the transcript counts show high overdispersion relative to Poisson even for technical replicates. In other words, RTA produces overdispersion even when the transcript proportions are constant, representing a failure in the underlying distributional assumptions for these packages.

Meanwhile, the *edgeR* and *limma* packages have also historically provided functions for DEU analysis that could be repurposed for DTU, including *diffSplice* in *limma* and *diffSpliceDGE* in *edgeR. diffSpliceDGE* fits negative binomial generalized linear models to the counts for each transcript and tests for differential usage via contrasts between the transcript-level log-fold-changes for each gene. *limma*’s *diffSplice* is similar but with normal linear models. This article presents recent improvements to both functions, and demonstrates that they provide fast and accurate tools for DTU when combined with divided transcript counts. Input for *limma*’s *diffSplice* is now prepared by *voomLmFit*, which improves on the older voom pipeline by adjusting the residual variance estimates for the loss of effective degrees of freedom (df) when there are experimental groups with all zero counts. *diffSplice* itself has been revised to accommodate the unequal residual df that now arise. *diffSpliceDGE* has been completely rewritten to provide more rigorous error rate control, to accommodate completely general experimental designs, and to make full use of the enhanced quasi-likelihood functionality of *edgeR* v4 [19]. In the Bioconductor 3.21 release of April 2025, *diffSplice* was converted to a be generic function that now handles DEU and DTU analyses for both *limma* and *edgeR* in a unified manner. The original non-generic function is now the *diffSplice* method for *limma* fitted model objects and the rewritten *diffSpliceDGE* is now the *diffSplice* method for *edgeR* fitted model objects produced by *glmQLFit*. The older *diffSpliceDGE* function is retained for backward compatibility, but is considered to be deprecated. The new DTU pipeline consists of *catchSalmon* to read Salmon quantifications and estimate RTA dispersions, followed by *voomLmFit* or *glmQLFit* to fit transcript-wise linear models to the divided counts, and then *diffSplice* for DTU. The *catchSalmon* function can be replaced with *catchRSEM* or *catchKallisto* if *RSEM* or *kallisto* are used instead of *Salmon*, but this article will focus on *Salmon* because we have previously found it to have good performance [6].

This article presents the statistical theory behind the revised *diffSplice* methods and benchmarks them against the other DTU methods described previously. An extensive simulation study shows the *edgeR* and *limma diffSplice* pipelines to be the best two performers, demonstrating greater statistical power than the other methods while also providing rigorous error rate control, controlling the FDR below the nominal rate even when the sample sizes are small. Comparison is also made with the *BANDITS* package, which tests for DTU directly from equivalence class counts [18]. The *diffSplice* methods are much faster than any of the competing methods, especially for larger sample sizes. They are also fully flexible, supporting completely general experimental designs with any number of covariates or batch effects. They allow DTU to be detected either between experimental conditions or as a function of a continuous predictor.

Given the effectiveness of divided counts for *limma* and *edgeR* analyses, we tested whether divided counts would improve the other methods as well. Divided counts were found to improve *DEXSeq*, bringing it closer to *edgeR* and *limma* performance, while the performance of *DRIMSeq* and *satuRn* did not improve.

The article finishes with a case study, using *diffSplice* to demonstrate the role of DTU in the lineage commitment of mouse mammary gland epithelial cells.

## Materials and methods

### User interface

The *limma* and *edgeR* DTU pipelines operate on transcript quantifications and technical replicates output by *Salmon, kallisto* or *RSEM*. The pipelines consist of three steps. First, the transcript quantifications are read by *catchSalmon, catchKallisto*, or *catchRSEM*, and the transcript counts are divided by the transcript RTA dispersions [5, 6]. We recommend filtering of low expressed transcripts with *filterByExpr* and library size normalization with *normLibSizes* at this stage. Second, the divided counts are input to either *voomLmFit* or *glmQLFit*, together with a design matrix describing the experimental setup. Finally, DTU is evaluated with the *diffSplice* and *topSplice* functions. Optionally, the fitted values from *voomLmFit* or *glmQLFit* can be converted to transcripts-per-million (TPM) and expression proportions with the *tpm* and *tpmProp* functions.

In *limma* 3.64.0 and *edgeR* 4.6.0, *limma*’s *diffSplice* function was converted to be an S3 generic, with methods defined for *limma* and *edgeR* fitted model objects. This unification means that the *limma* and *edgeR* DTU pipelines are now operationally the same, except for the use of either *voomLmFit* or *glmQLFit* at the second step. *diffSplice* stores gene and transcript level tests so that *topSplice* can extract whichever tests are required. The main user difference between the *edgeR* and *limma* methods is that the *edgeR* method evaluates DTU for one coefficient or contrast at a time, as specified by the call to *diffSplice*, whereas the *limma* method evaluates DTU for all the coefficients in the linear model at once. The detailed mathematical basis of the two pipelines is described in the following four sections.

### RTA-divided counts

The divided-count approach was introduced by Baldoni *et al*. [5, 6], *and is an essential component of the limma* and *edgeR* pipelines for DTE and DTU. Consider an RNA-seq experiment consisting of *n* samples and transcriptome annotation with *T* transcripts from *G* genes. Write *y*_*gti*_ for the read count for transcript *t* of gene *g* in sample *i*, as estimated by *Salmon*, and let 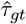 be the RTA-dispersion for that transcript, as estimated by *catchSalmon* [6]. Then the divided counts 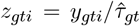 can be treated as approximately negative binomial distributed counts [5, 6, 19]. The divided counts *z*_*gti*_ are input into the *limma* or *edgeR* analysis pipelines as if they were ordinary read counts. The pipelines can be applied similarly to *kallisto* or *RSEM* quantifications via the *catchKallisto* or *catchRSEM* functions, although *Salmon* quantifications are used in this article.

### voomLmFit and glmQLFit

The functions *voomLmFit* or *glmQLFit* fit transcript-wise linear models to the divided counts. *voomLmFit* fits linear models to log2-counts-per-million, and is a further development of the popular limma-voom approach [20]. Unlike the original voom function, it reduces the residual df to avoid underestimation of the residual variance when a transcript has entirely zero counts for any treatment group [28]. It also automates the estimation of sample quality weights [29] and the incorporation of random effect blocks [30].

*glmQLFit* fits quasi negative binomial generalized linear models to the transcript counts, with bias adjusted deviances and empirical Bayes moderation of the quasi-dispersions [19]. Like *voomLmFit, glmQLFit* also adjusts the residual df for zero counts, but in a continuous way, so that fitted values that are very small but positive also generate reduced df.

*voomLmFit* and *glmQLFit* accept design matrices, which define the experimental conditions and the associated linear model to be estimated [31], and return estimated linear model coefficients 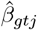 for each column *j* of the design matrix.

Transcript-level RNA-seq counts tend to be sparser than gene-level counts because of the larger number of transcripts compared to genes, so the robustness to small counts offered by *voomLmFit* and *glmQLFit* is important. *voomLmFit* and *glmQLFit* both generate transcript-specific residual df, and make use of *limma*’s *fitFDistUnequalDF1* function for empirical Bayes hyperparameter estimation.

### limma’s diffSplice method

We now describe the statistical theory of the *diffSplice* method applied to *limma* linear models. The DTU analysis focuses on a particular coefficient *j* of the linear model, which typically corresponds to the log2-fold-change (logFC) of interest. The method is extremely general and can be applied to any meaningful coefficient or contrast, including a coefficient for a continuous covariate or for a paired-samples treatment. The simulations reported in this article used simple two-column design matrices where the second column represents the loge-fold-change in expression between the two groups.

Let *T*_*g*_ be the number of transcripts for gene *g*, so that *T* = Σ*T*_*g*_. DTU tests are performed on genes expressing *T*_*g*_ *>* 1 transcripts and are equivalent to testing whether the transcript-specific coefficients 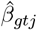 differ among transcripts of the same gene. Rejecting the null hypothesis of equality implies that the relative expression of the transcripts must differ between conditions or with respect to the relevant linear model covariate.

For notational simplicity, we will drop the *j* subscript for the rest of this section, with the understanding that all the tests relate to coefficient *j* in the linear model.

The fundamental distributional assumptions made by *diffSplice* are that the transcript-wise coefficients 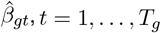, are statistically independent and that they share the same gene-wise residual error variance, 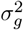. *diffSplice* assumes that

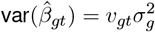

where *v*_*gt*_ is the unscaled variance for the coefficient arising from the experimental design and from the voom precision weights for that transcript. Write *u*_*gt*_ = 1*/v*_*gt*_ for the unscaled precision. The genewise coefficient averaged over the transcripts is

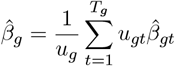

where *u*_*g*_ = Σ_*t*_ *u*_*gt*_ is the total precision summed over all the transcripts for the gene. Also write

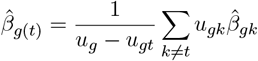

for the coefficient averaged over all the other transcripts for the gene, excluding transcript *t*. Note that 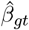 and 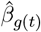 are statistically independent. The transcript-level tests for DTU are based on the differences 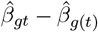 for each transcript *t* in turn. It can be shown that

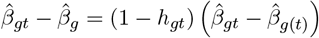

where *h*_*gt*_ = *u*_*gt*_*/u*_*g*_ is the leverage (or relative weight) given to transcript *t* in the genewise coefficient. It can also be shown that

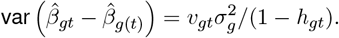

It follows therefore that

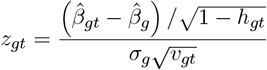

has a standard normal distribution under the null hypothesis.

Let

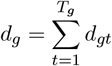

be the total residual df over all the transcripts for gene *g*, and let

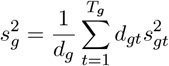

be the pooled residual variance for gene *g. limma*’s empirical Bayes variance estimation is applied to the gene-wise 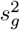, resulting in prior df *d*_0*g*_ and posterior variance estimators 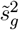 [25, 26]. It follows that the empirical Bayes moderated t-statistic

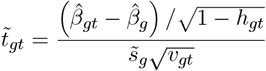

follows a *t*-distribution on *d*_0*g*_ + *dg* df under the null hypothesis. Also define the moderated F-statistic

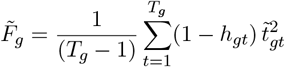

It can be shown that 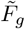 follows a *F* -distribution on *T*_*g*_ − 1 and *d*_0*g*_ + *d*_*g*_ df under the null hypothesis. The moderated F-statistic can be used to test the overall null hypothesis of no DTU for gene *g*, whereas the moderated t-statistic can be used to test for differential usage of a particular transcript within the gene. The two tests are equivalent in the sense that rejecting any of the transcript-wise null hypotheses for a gene is equivalent to rejecting the overall null hypothesis.

Genewise p-values for DTU can be obtained from the F-statistic and transcript-specific p-values can be obtained from the moderated t-statistics. *diffSplice* also offers gene-wise p-values obtained from applying Simes adjustment to the transcript-level p-values for each gene [32]. The F-statistic and Simes p-values are identical for genes with just two transcripts but differ for genes with many transcripts, with the Simes p-values tending to be smaller than the F-statistic p-values when one of the 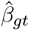 coefficients is an outlier compared to the others for the same gene. Gene-wise p-values can then be adjusted for multiple testing and genes can be ranked by evidence for differential usage with the *topSplice* function of the *limma* package.

### edgeR’s diffSplice method

The *edgeR* method follows the same format as that for *limma*, but uses quasi F-statistics obtained from deviance differences. The divided counts are assumed to follow a quadratic-mean variance relationship

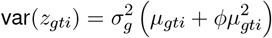

where *µ*_*gti*_ is the expected count and *ϕ* is a global negative binomial dispersion estimated by *glmQLFit*.

The quasi-dispersions 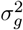 are estimated in exactly the same way as the residual variances in the *limma* pipeline, except that residual deviances replace residual sums of squares. If *d*_*gt*_ is the bias-adjusted residual df and 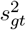 is the biased-adjusted mean deviance for transcript *t*, then

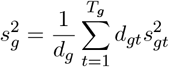

is the pooled mean deviance for gene *g. limma*’s empirical Bayes variance estimation is applied to the 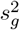, resulting in posterior quasi-dispersion estimators 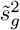.

The gene-level quasi F-statistic is

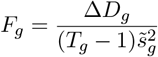

where Δ*D*_*g*_ is the deviance difference between the full model and the null model with all the *β*_*gt*_ equal, *t* = 1, …, *T*_*g*_. The transcript-level quasi F-statistic is

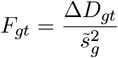

where Δ*D*_*gt*_ is the deviance difference between the null model and the one-removed null model in which all the 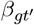 are equal except for the transcript of interest, i.e., for *t*^*′*^ ≠ *t*. The transcript-level F-statistic is on 1 and *d*_0*g*_ + *d*_*g*_ df, and can be transformed to a t-statistic by

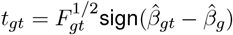

where 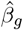 is the consensus value for the linear model coefficient over all the transcripts for that gene.

For genes with a very large number of transcripts (*>* 10 by default), a fast approximation to *F*_*gt*_ is used. The approximation replaces Δ*D*_*gt*_ with the deviance difference between setting 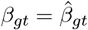 vs setting 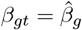.

### Relationship to TPMs

We indicate briefly here how the *diffSplice* transcript-wise linear models are mathematically equivalent to a DTU model in terms of expression proportions. The *limma* and *edgeR* analyses do not require the computation of transcripts per million (TPM) expression values, but TPM values can be recovered from the *edgeR* fitted values after rescaling by the RTA dispersions and the transcript lengths. For simplicity, we consider a simple experimental design with two conditions, A and B, and a gene with two transcripts. Also assume a standard R factor parametrization, so that *β*_*gt*1_ represents baseline expression in condition A and *β*_*gt*2_ is the logFC in condition B. Note that *edgeR* coefficients are computed and stored internally on the natural log-scale, although user results are reported on the log2-scale by functions such as *topTags* or *topSplice. edgeR* linear models also include the log library size as an offset. Table 1 gives the TPM values for the two transcripts in the two conditions where

**Table 1:** TPM in terms of *edgeR* linear model coefficients for a gene with two transcripts and an experiment with two conditions. The factors in the table are defined in the manuscript text.

|  | Condition A | Condition B |
| --- | --- | --- |
| Transcript 1 | $\alpha_1$ | $\alpha_1\gamma$ |
| Transcript 2 | $\alpha_2$ | $\alpha_2\gamma\delta$ |

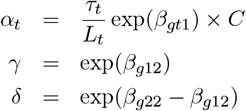

Here *τ*_*t*_ is the RTA dispersion for transcript *t, L*_*t*_ is the effective transcript length, and *C* is an overall scaling factor constant for all samples and transcripts to ensure that total TPM is approximately one million per sample. The terms *α*_1_ and *α*_2_ are the baseline TPM expression levels in condition A, *γ* is the fold-change of transcript 1 in condition B, and *δ* is the transcript × condition interaction, If the two transcripts have the same logFC, then *δ* = 1 and the proportion of the gene’s TPM expression arising from transcript 2 will be equal to *α*_2_*/*(*α*_1_ + *α*_2_) in both conditions. This formulation shows explicitly the equivalence of the *diffSplice* null hypothesis *β*_*g*12_ = *β*_*g*22_ to DTU in terms of TPMs.

In *edgeR*, TPM values can be extracted from the fitted model object created by *glmQLFit* using the *tpm* function, and these can be converted to expression proportions for each gene using the *tpmProp* function. The *edgeR* TPM values incorporate effective library sizes as estimated by *normLibSizes*, which allow for the possibility that total RNA production may differ from sample to sample [33]. The total sum of TPM values will therefore differ from sample to sample, with the total sum being inversely related to the normalization factor for that sample. The *tpm* function sets the geometric mean of the column totals to be equal to one million. The TPMs preserve the same fold-changes as estimated by *glmQLFit*.

### RNA-seq profiling of mouse mammary gland epithelial cell populations

RNA-seq data was obtained from NCBI Gene Expression Omnibus (GEO) series GSE227748, which provides transcriptome profiles of freshly sorted epithelial cell subpopulations from the mammary glands of adult female FVB/N mice. Myoepithelial (basal), luminal progenitor (LP) and mature luminal (ML) cells were sorted from three mice, providing three biological replicates of each cell population. Libraries were sequenced using an Illumina NextSeq 500 system, producing 35–80 million 80 bp read-pairs per sample. *Salmon* was run on the FASTQ files with default options using the decoy-aware transcriptome index generated from M35 Gencode transcript annotation and the mm39 mouse genome. 100 Gibbs resamples were generated for every sample. *catchSalmon* was used to import *Salmon*’s quantifications and to estimate the RTA overdispersion parameters. Divided transcript counts were computed, low-expressed transcripts were filtered by *filterByExpr*, genes other than protein-coding and lncRNA were removed, library sizes were normalized by *normLibSizes* and DTU was assessed using the *limma* and *edgeR diffSplice* methods with *robust=TRUE*.

### Simulated datasets

The mouse mammary gland data was used to select a reference set of genes and transcripts from which sequence reads would be simulated. The reference set of genes and transcripts was created as follows. Gene-level counts were computed by adding up transcript-level expected counts obtained from *Salmon*. Only genes annotated to the mouse autosome or sex chromosomes, and classified as protein-coding or lncRNA were considered. Genes with duplicated transcript sequences were discarded. The nucleotide fraction attributable to each gene in each sample was estimated by the Good-Turing algorithm [34], implemented in *edgeR*’s *goodTuringProportions* function. Genes with nucleotide fractions greater than 10^−6^ in at least six of the nine RNA-seq libraries were retained, along with their protein-coding and lncRNA transcripts, for inclusion in the reference set. The resulting reference set comprised 12,715 genes and 41,554 associated transcripts.

Transcript-level dispersion parameters (squared BCV) were generated from an inverse chisquared distribution with scale equal to 0.25^2^ and df equal to 40, i.e., 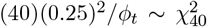. where *ϕ*_*t*_ is the dispersion parameter. This choice of 0.25 for the prior BCV matches the biological variability and heterogeneity that we typically observe in our own in-house RNA-seq experiments using genetically identical laboratory mice [20]. Sample-wise TPMs for each transcript were generated from a gamma distribution with squared CV equal to *ϕ*_*t*_ and baseline expected value generated from the distribution of Good-Turing proportions from the reference dataset.

For each simulated dataset, we randomly selected a set of 4,500 multi-transcript genes to contain at least one differentially expressed transcript. Genes from this set were randomly selected to exhibit either differential usage alone (*n* = 1, 500), a combination of differential usage and differential expression (*n* = 1, 500), or differential expression alone without differential usage (*n* = 1, 500). Genes exhibiting differential usage (DTU-only) alone had the expected TPM of two of their transcripts adjusted and then swapped in one of the groups. This process ensured a two-fold-change for the two selected transcripts while keeping the total expression of the gene constant in both groups. For such genes, only the two differentially expressed transcripts were considered to be differentially used. Differentially used and expressed genes (DGE/DTU) had the expected TPM of one of their transcripts adjusted with a two-fold-change in one of the groups. For such genes, all their transcripts were considered to be differentially used because, when the adjusted transcript increases expression, all other transcripts of the same gene must necessarily contribute a lower proportion of the total expression of that gene. Genes selected to be differentially expressed alone (DGE-only) had the expected TPM of all of their transcripts adjusted with a two-fold-change in one of the groups. For these genes, no transcripts were considered to be differentially used. Differentially expressed genes were randomly assigned to be up or down regulated, so that the simulated datasets had a balanced number of up and down regulated transcripts overall.

Null simulations without any real differential expression or usage between groups were also generated to assess type I error rate control. For the null simulations, each transcript had the same expected TPM across samples.

The TPMs for each transcript in each sample were then input to the *simReads* function of the *Rsubread* package [35] to generate a FASTQ file of sequence reads for each sample. Paired-end reads with 100 base pairs (bp) were generated in scenarios that varied with respect to the number of biological replicates per group (three, five, or ten) with unbalanced library sizes (either 25 or 100 million reads per sample). For each scenario, 20 simulated experiments with RNA-seq libraries from two groups were generated.

The simulated data sets were quantified with *Salmon* with a decoy-aware mapping-based indexed transcriptome generated from the mouse mm39 reference genome with k-mers of length 31 and the complete mouse Gencode transcriptome annotation M35. 100 Gibbs resamples were generated for every sample.

### Software tools included in benchmarking

The performance of the *limma* and *edgeR diffSplice* methods was compared to that of *DEXSeq, DRIMSeq*, and *satuRn* on the simulated datasets. Methods were evaluated with respect to type I error rate control, false discovery rate (FDR), power, and computational speed. All methods are available as Bioconductor packages for the R programming language [36]. The *limma* and *edgeR* methods were implemented as described above with default settings for all functions. *DEXSeq, DRIMSeq*, and *satuRn* were run initially on raw transcript counts, as recommended in their original publications, and then also on the divided transcripts counts. For all methods but *DRIMSeq*, low expressed transcripts were filtered out with *edgeR*’s *filterByExpr* function with default settings. For pipelines using divided counts, the counts were divided before running *filterByExpr*. For *DRIMSeq*, its own *dmFilter* function was used to select transcripts with a minimum count of 10 in at least *n* samples, where *n* is the group sample size. Empirical p-values were used to assess DTU with *satuRn*, as recommended by the authors of that package [16]. To standardize timings, all the above analyses were conducted on WEHI’s high performance Linux system using a single core.

A second set of DTU analyses was conducted to evaluate the performance of *edgeR* and *limma* under a less stringent filtering criteria, retaining transcripts with at least 1 count in at least half the samples (equivalent to *filterByExpr* with *min*.*count=1* and *min*.*total*.*count=5*).

The *BANDITS* package was benchmarked separately, because it requires different input to the other tools and because it could not be run successfully with a single core. *BANDITS* requires transcript counts from *Salmon* for transcript filtering and hyperparameter estimation and equivalence class counts for the DTU analysis, but doesn’t use technical replicates because it estimates the uncertainty arising from RTA directly from likelihood modelling of the equivalence class counts [18]. Transcripts were filtered using *filter transcripts*, as recommended in the package documentation. *BANDITS* was run using 10 cores on five simulated datasets for each simulation scenario.

## Results

### Divided counts restore the biological mean-variance relationship

Figure 2 of Baldoni et al. [5] displayed *edgeR* dispersion plots for raw and divided counts, showing that the mean-variance relationship associated with biological variation in RNA-seq counts in disrupted in the raw transcript counts from *Salmon* but restored in the divided counts. The behaviour was very similar for both simulated and real RNA-seq data. Supplementary Figure S1 shows the same behaviour for the current simulations and for the mouse mammary data considered in this article.

### limma and edgeR control the type 1 error rate correctly

A fundamental requirement of any statistical test is that it should yield valid p-values when the null hypothesis is true, i.e., that it should control the type I error rate at or below the nominal rate. Ideally, a test that balances sensitivity with error rate control will produce null p-values that are uniformly distributed between 0 and 1, ensuring that the type I error rate is always equal to the nominal level. We started therefore by examining the p-value distributions generated by each of the DTU methods in the null simulations, with no differential usage or differential expression between groups. Figure 1 presents density histograms of raw p-values from all the methods for the null simulation scenario with five samples per group. *limma* and *edgeR* are the only methods that yield uniform p-value distributions, with both transcript-level t-tests and gene-level F-tests giving p-value histograms that are very close to uniform. Simes-adjusted p-values from *limma* and *edgeR* show slightly U-shaped histograms, although the density near zero remains below the nominal level. *DEXSeq* and *satuRn* show conservative distributions, somewhat depleted for small p-values, while *DRIMSeq* produces spikes at 0 and 1, indicating excessive numbers of both very large and very small p-values. The results were similar with *n* = 3 or *n* = 10 samples per group, except that *satuRn* become noticeably less conservative at the largest sample size (Supplementary Figures S2–S3).

**Figure 1:**
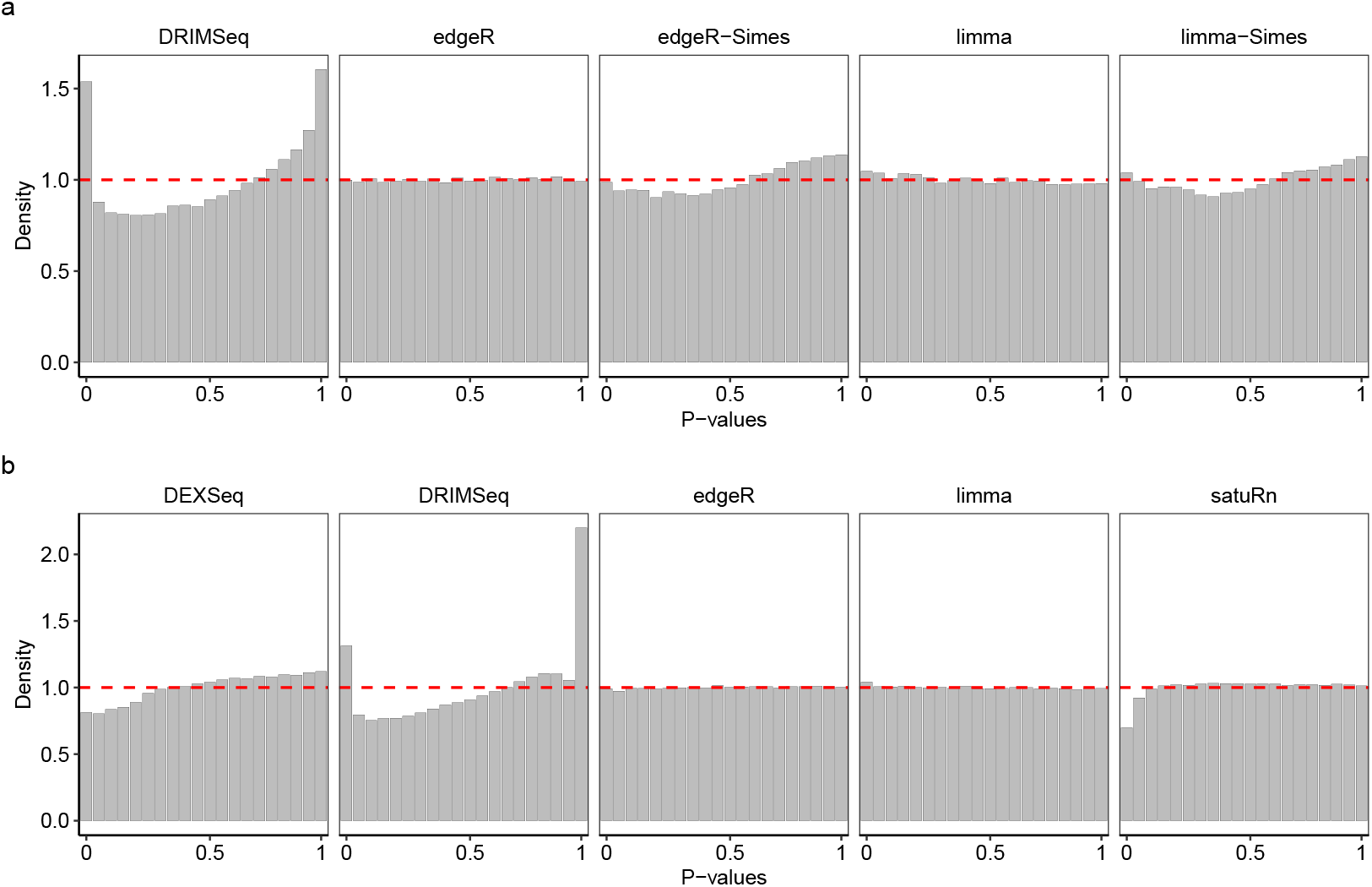
Density histograms of p-values in the null simulation with five samples per group. Dashed line indicates the expected uniform distribution. Panels (a) and (b) show results at the gene and transcripts levels, respectively. Methods DEXSeq and satuRn do not report gene-level raw p-values and, therefore, are omitted from panel (a). Raw counts were used by *DEXSeq, DRIMSeq*, and *satuRn*. Results are averaged over 20 simulated datasets.

Supplementary Figure S4 shows the actual type I error rates at the nominal 5% level, confirming the qualitative observations made from the density histograms. *limma* and *edgeR* give type I error rates very close to the nominal level, whereas *DEXSeq* and *satuRn* are conservative and *DRIMSeq* is liberal.

Here and in the next two sections, we compare each of the methods as they were originally proposed, meaning that *limma* and *edgeR* use divided counts while all the other methods use raw transcripts counts. Later we explore whether divided counts can improve the other methods as well.

### edgeR and limma give the best gene and transcript rankings

Next we compared the ability of the different methods to rank genes and transcripts in terms of evidence for DTU when true DTU is present. Figure 2 shows cumulative false discovery rates (FDRs) when genes or transcripts are ranked by p-value for each DTU method. At both the gene and transcript levels, and for any sample size, *edgeR* selected the smallest number of false discoveries for any chosen number of features, followed closely by *limma*. The F-test and Simes-adjusted p-values performed similarly. *DEXSeq* was next in performance, but with considerably higher FDRs than *edgeR* or *limma. satuRn* gave poor results with *n* = 3 samples per group but improved noticeably at the larger samples sizes.

**Figure 2:**
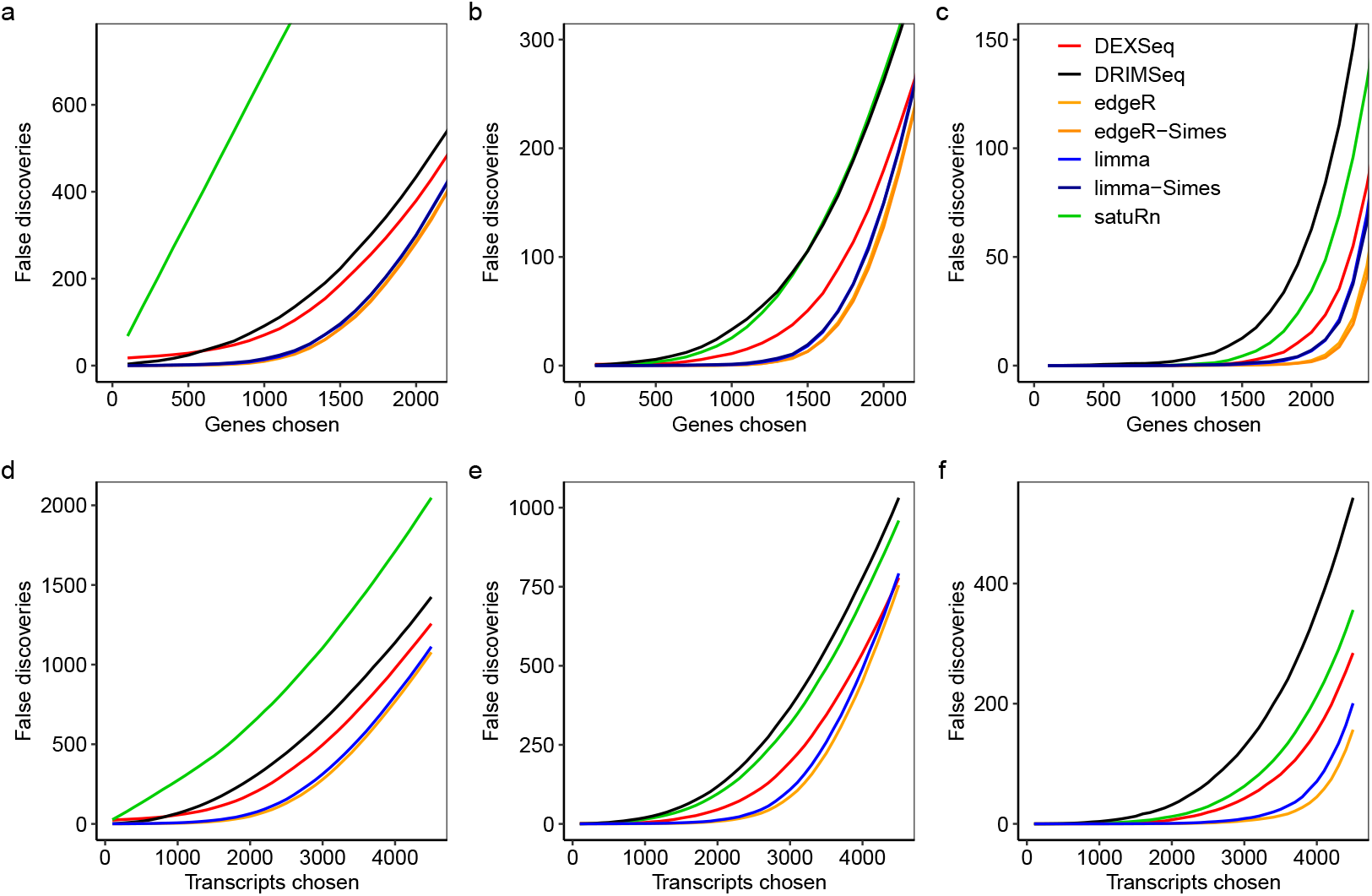
Cumulative false discoveries in ranked lists. The number of false discoveries is plotted for each method versus the number of genes or transcripts selected as differentially used. Panels (a)–(c) show gene-level results with three, five, and ten samples per group, respectively. Panels (d)–(f) show transcriptlevel results with three, five, and ten samples per group, respectively. For *DEXSeq* and *satuRn*, genes were ranked according to their adjusted p-values, as those methods do not provide unadjusted p-values by default. Raw counts were used by *DEXSeq, DRIMSeq*, and *satuRn*. Results are averaged over 20 simulations.

### edgeR and limma maximise power while maintaining FDR control

Finally, we examined the combined effect of error rate control and feature ranking at common FDR cutoff levels by counting the numbers of true and false discoveries. Figure 3 shows the observed number of true and false discoveries at an FDR cutoff of 0.05. *edgeR* controlled the FDR below nominal level in all scenarios, while also yielding more true discoveries than any other method except *DRIMSeq*, which produced unacceptably high FDRs. *limma* was nearly as good as *edgeR*, although was slightly liberal at the transcript level with the larger sample sizes. *DEXSeq* was somewhat less powerful than *edgeR* or *limma*, and also failed to control the FDR correctly in four out the six scenarios. *satuRn* was the least powerful method, making very few or no discoveries at all with sample sizes less than 10.

**Figure 3:**
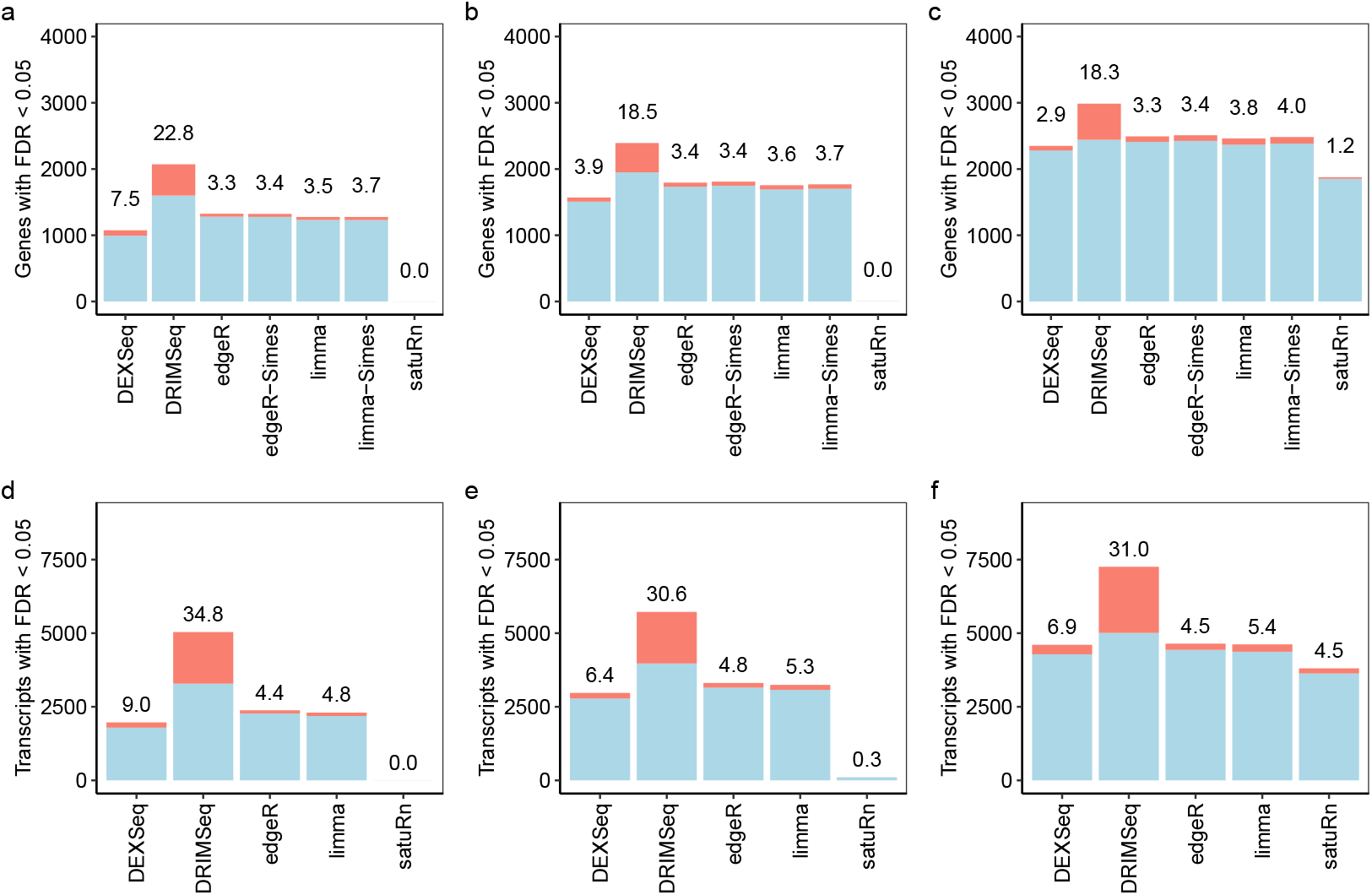
Numbers of true and false discoveries at 5% FDR. Stacked barplots show the number of true (blue) and false (red) positive differentially used genes and transcripts at nominal 5% FDR. The observed FDR is shown as a percentage over each bar. Panels (a)–(c) show results at the gene-level with three, five, and ten samples per group, respectively. Panels (d)–(f) show results at the transcript-level with three, five, and ten samples per group, respectively. *satuRn* failed to detect differentially used genes or transcripts with three samples per group. Raw counts were used by *DEXSeq, DRIMSeq*, and *satuRn*. Results are averaged over 20 simulated datasets.

Supplementary Figure S5 shows true and false discoveries at FDR cutoff 0.01. At this more stringent cutoff, the results are qualitatively similar to 0.05 but more striking. Again, *limma* and *edgeR* control the FDR at close to the correct level, but now they have more power even than DRIMSeq at the gene-level with *n* = 10. *DEXSeq* is very liberal when *n* = 3, with FDR six times the nominal level, and is liberal at the transcript level for all sample sizes. *DRIMSeq* remains too liberal and *satuRn* too conservative.

Supplementary Figure S6 plots, for each method, the observed power versus the observed FDR for nominal FDR cutoffs ranging from 0.01 to 0.20. This confirms previous results, with *edgeR* and *limma* at the top left of each of the panels.

### Relative power of Simes vs F-tests depends on the DTU configuration

*diffSplice* provides two summary p-values for each gene, one derived from an F-test and the other from Simes adjustment of the transcript-level t-tests for the same gene. The motivation for providing two different p-values is that they may have different statistical power in different DTU contexts. Our simulation included two different DTU scenarios, with either one or two differentially expressed transcripts per gene. In the latter case, the two transcripts were differentially expressed in opposite directions, switching expression levels with one another in the two experimental conditions. Supplementary Figure S7 plots the F-test vs Simes p-values for the two scenarios for a representative simulated dataset with ten samples per group. The F-test p-values tend to be smaller than the Simes p-values when there are two differentially expressed transcripts (Supplementary Figure S7a), whereas the Simes p-values tend to be smaller when there is only one differentially expressed transcript. This confirms our expectation that Simes test should be more powerful when there is a single stand-out t-statistic, while the F-test should be more powerful in more complex situations involving multiple transcripts. The difference between the two methods is more marked for genes with more transcripts. The F-test and Simes p-values are identical for genes with only two transcripts in total, but become more different as the total number of transcripts increases. As expected, the p-values are smaller overall in panel (a) than in panel (b) of Supplementary Figure S7, showing that it is easier overall to detect DTU when there are multiple transcripts changing in opposite directions.

### Divided counts improve DEXSeq but not DRIMSeq or satuRn

Given the success of divided counts in the *edgeR* and *limma* pipelines, we investigated how the use of divided counts instead of raw transcript counts would affect *DEXSeq, DRIMSeq*, and *satuRn*. We repeated the analysis of the simulated data using divided counts for all benchmarked methods (Supplementary Figures S8–S14). The use of divided counts noticeably improved the performance of *DEXSeq*, although *edgeR* and *limma* remained the best two methods. For *DEXSeq*, the conservative depletion of small p-values previously seen in Figure 1 and Supplementary Figure S3 was replaced by some anti-conservative enrichment of small p-values for the *n* = 5 and *n* = 10 scenarios (Supplementary Figure S9). With divided counts, *DEXSeq* was able to achieve good FDR control at the gene-level even for *n* = 3, although *edgeR* continued to have higher power as well as lower FDRs (Supplementary Figures S11–S12). At the transcript-level, *DEXSeq* became more powerful but was now overly liberal at all sample sizes and FDR cutoffs. The performance of *satuRn* becomes worse in terms of cumulative FDR (Supplementary Figure S13). The performance of *DRIMSeq* is improved, but only slightly, and its FDR remains unacceptably high at all nominal levels (Supplementary Figure S14). We conclude that the divided count approach aids the performance of all the methods that assume a negative binomial distribution for transcript-level counts, but gives no substantial improvement for methods that make binomial or multinomial assumptions.

### Comparison with *BANDITS*

Benchmarking was also conducted of the *BANDITS* package, as a way to compare the divided count pipelines with an approach that infers DTU directly from the equivalence class counts. *BANDITS* produces null p-value histograms that overall are strongly skewed towards large p-values, but which also include a substantial spike at very small p-values, many of which are exactly zero (Supplementary Figure S15). This p-value pattern, enriched for both large and very small p-values, is associated with statistical power that is lower than *edgeR* or *limma* but also with inflated FDRs (Supplementary Figure S16). *BANDITS* was found to have lower power but higher FDR than *edgeR* or *limma* in all scenarios: at both gene and transcript levels, for all sample sizes, and at all significance cutoffs. *BANDITS* did control the 5% FDR level correctly at the gene level, but was very liberal at the transcript level and for more stringent significance cutoffs. *DEXSeq* with divided counts was also uniformly better than *BANDITS* at the gene level, although the comparison between the two was mixed at the transcript level, where both packages were very liberal.

### limma and edgeR are faster than other DTU methods

We recorded the computation time required by each of the DTU methods for the simulation study (Figure 4). *DRIMSeq* was by far the most time-consuming of the transcript-count methods, followed by *DEXSeq* in scenarios with large sample sizes. *satuRn* scaled well computationally as the number of replicate samples increased, being slower than *DEXSeq* for *n* = 3 but much faster for *n* = 10. *limma* and *edgeR* were by far the fastest methods, taking only about 20 seconds for each analysis even for the largest sample sizes. *limma* evaluates DTU for all coefficients in the linear model simultaneously, so it will be especially efficient for large-scale experiments in which multiple treatment comparisons need to be made and DTU is assessed for each one.

**Figure 4:**
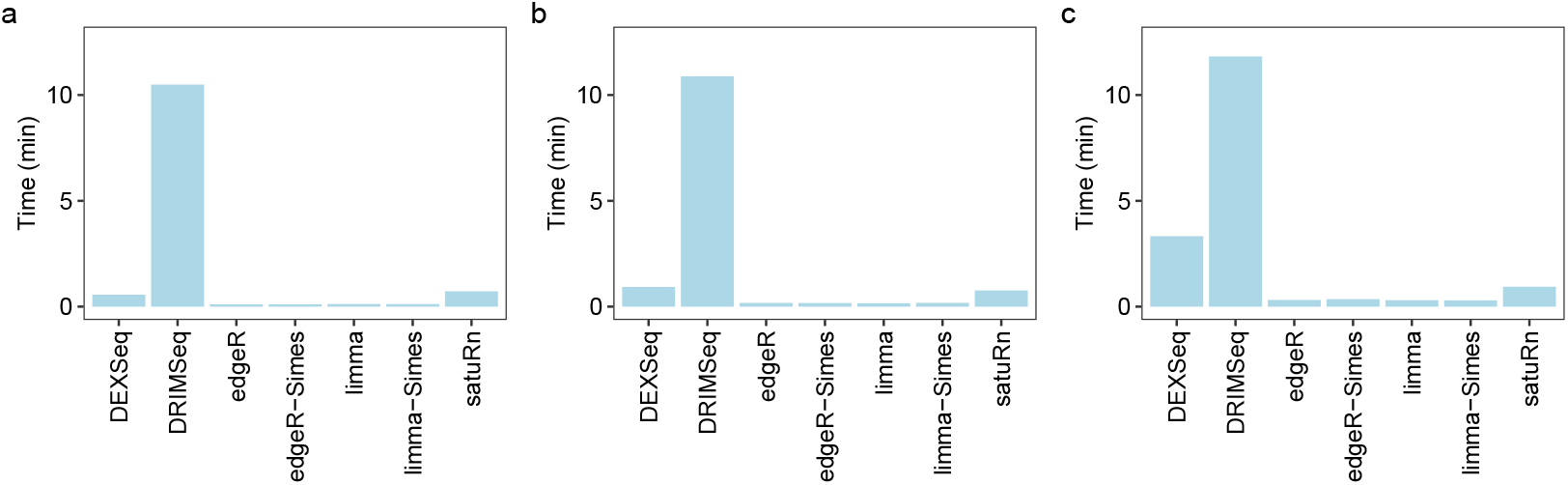
Computation time. Bars show time in minutes required for a DTU analysis by the different methods. Panels (a)–(c) show results with three, five, and ten samples per group, respectively. *BANDITS* is not shown on the plot because it was much slower than the other methods and required more computing cores, taking 16.8, 26.3 and 81.3 minutes respectively for three, five and ten samples per group. Analyses were run on a Linux system using 10 cores for *BANDITS* and a single core for the other methods. Results are averaged over 5 simulations for *BANDITS* and 20 simulations for the other methods.

*BANDITS* was much slower than the transcript count methods and also required the use of multiple cores (Figure 4 legend). A *BANDITS* analysis can recover some time relative to the other methods by omitting resampling when running *Salmon*, but Gibbs sampling will typically take only a few minutes in total [6, Table 1].

### limma and edgeR are robust to transcript filtering

Finally, we assessed the performance of the *limma* and *edgeR* DTU pipelines under less stringent filtering criteria. Specifically, we re-analyzed the simulated datasets using *filterByExpr* with *min*.*count=1* and *min*.*total*.*count=5. edgeR* continued to control the FDR correctly under these more lenient criteria, and detected more true positives for *n* = 5 and *n* = 10 but not for *n* = 3 (Supplementary Tables S1–S2). This suggests that more lenient filtering could be beneficial for the larger sample sizes. The FDR is relatively high though for the extra discoveries. For instance, for *edgeR* and *n* = 10, lenient filter gave 151.55 more true positive DTU genes on average at a cost of 21.90 more false positives, corresponding to a FDR of 17% among the extra discoveries. With lenient filtering, *limma* became slightly liberal at the transcript level tests with *n* = 3 (Supplementary Table S2), although the FDR control for low counts could be improved by setting the *robust=TRUE* when running *diffSplice* (data not shown).

### Differential transcript usage in the steady-state adult mouse mammary gland

The transcriptomic profile of the mouse mammary stem cell lineage has been extensively studied in the literature. The steady-state mammary gland of adult mice comprises three major cell types: basal, luminal progenitor (LP), and mature luminal (ML) cells. A number of genes, such as *Foxp1, Ezh2*, and *Asap1*, have been shown to be essential for the morphogenesis of the mammary gland and to regulate cell-specific differentiation programs [37, 38, 39]. Importantly, recent work has identified several genes exhibiting alternative transcriptional start-site usage across the stem cell lineage, including genes known to be indispensable for mammary gland development, such as *Foxp1* [7]. Altogether, these results suggest that differential usage of transcripts might be an important mechanism of mammary stem cell lineage differentiation. To investigate this phenomenon further, we performed a DTU analysis of the Illumina paired-end RNA-seq data from the major cell types comprising the steady-state mouse mammary gland as reported by Milevskiy *et al*. [7]. The experiment includes nine RNA-seq samples from basal, LP, and ML cells, with three biological replicates per cell type.

The analysis presented here focuses on the basal versus LP cells comparison (Figure 5). We applied the *edgeR* pipeline to detect differential usage both at the gene-level with moderated quasi F-tests as well as at the transcript-level with moderated t-tests. We first considered transcript-level raw counts and observed a strong transcript-specific RTA overdispersion effect in the BCV of expressed transcripts (Figure 5a). The RTA overdispersion effect led the estimated BCV from transcripts associated with high RTA to systematically deviate from the typical trend that decreases smoothly with abundance commonly observed in gene-level read counts [20]. With RTA-divided transcript counts, the estimated dispersions reflect the true precision of effective counts, which in turn represent the biological variation of the RNA-seq experiment and indicate that RTA-divided transcript counts could be readily input into *edgeR*’s quasi negative binomial generalized linear modeling pipeline (Figure 5b). In contrast to RTA-divided counts, negative binomial dispersions estimated from raw counts resulted in systematically lower estimated gene-level prior df during the empirical Bayes moderation of estimated gene-wise quasi-dispersions (4.117 vs. 8.170, on average). The lower estimated prior df indicates that gene-wise quasi-dispersions estimates are less strongly moderated, resulting in an associated loss of statistical power to detect differences in usage with transcript-level raw counts.

**Figure 5:**
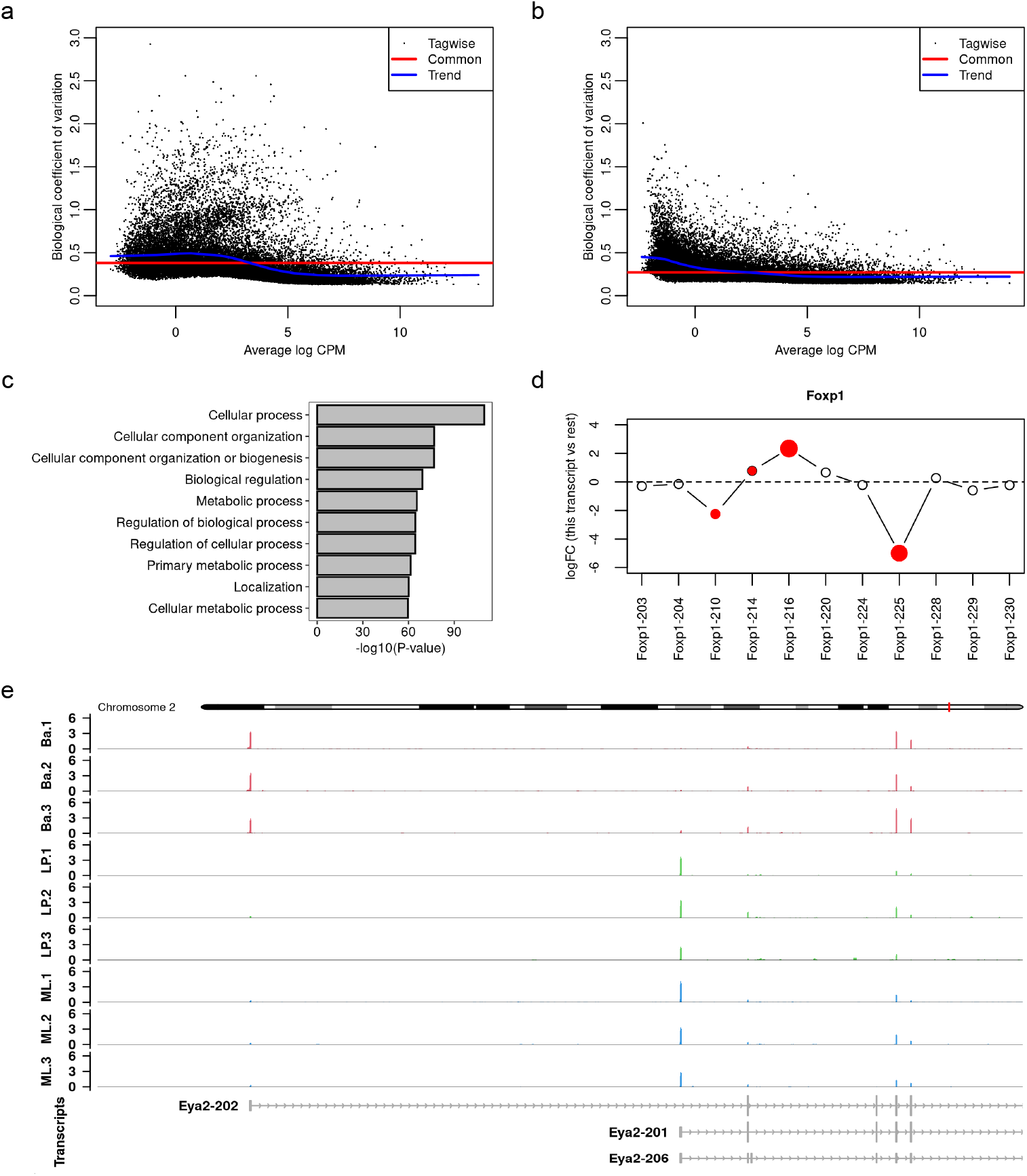
Panels (a)–(e) show the main results from the RNA-seq DTU analysis of the adult mouse mammary gland data. In (a), BCV plot using transcript raw counts. In (b), BCV plot using transcript divided counts. In (c), top 10 most significant over-represented gene ontology terms associated with biological processes from differentially used genes between basal and LP cells. In (d), differential splicing plot of relative log2 fold-changes (logFC) of expressed Foxp1 transcripts. The relative logFC is the difference between the transcript’s logFC and the overall logFC for the gene. The Foxp1 gene is differentially used between basal and LP cells (FDR = 3.3 × 10^−13^). Foxp1 transcripts 216, 225, 210, and 214 are differentially used with respect to other Foxp1 transcripts (FDR = 7.3 × 10^−8^, 2.9 × 10^−7^, 7.0 × 10^−3^, and 4.9 × 10^−2^, respectively). In (e), coverage profile of the differentially used gene Eya2 between basal and LP cells exhibiting cell-specific transcriptional start-site usage. Coverage profiles are defined as the number of reads overlapping each base, transformed into a count-per-million value based on effective library sizes. Coverage tracks include all expressed protein-coding and lncRNA transcripts based on the mouse Gencode annotation version M35. Nominal FDR of 0.05 in both gene- and transcript-level DTU analyses.

We identified a total of 1,256 differentially used genes between basal and LP cells using moderated genelevel quasi F-tests with FDR cutoff 0.05. Among these, we detected differential usage for several key genes previously reported in the literature as being involved in the regulation of mammary stem cell lineage differentiation. Notable examples include *Foxp1* (FDR= 3.3 × 10^−13^), *Ezh2* (FDR= 7.5 × 10^−3^), and *Asap1* (FDR= 2.6 × 10^−7^). The top most differentially spliced gene was *Tpm1* (FDR = 2.348 × 10^−38^), a gene responsible for stress fiber formation in human epithelial cells whose downregulation in breast tumors has been linked to the survival of neoplastic cells and tumor growth [8, 9]. We tested for over-representation of gene ontology terms associated with biological processes in the set of differentially spliced genes. Our analysis indicated that differentially spliced genes between basal and LP cells were mostly associated with cellular metabolic and regulatory processes (Figure 5c).

At the transcript-level, we have identified 2,478 differentially used transcripts between basal and LP cells using moderated transcript-level t-tests with FDR cutoff 0.05. Of particular interest was the detection of four differentially used transcripts from the *Foxp1* gene: the protein-coding transcripts *Foxp1-216* (relative logFC = 2.34, FDR = 7.3 × 10^−8^), *Foxp1-225* (relative logFC = −4.99, 2.924 × 10^−7^), *Foxp1-210* (relative logFC = −2.246, FDR = 7.0 × 10^−3^), and *Foxp1-214* (relative logFC = 0.78, FDR = 4.9 × 10^−2^) (Figure 5d). The relative logFC represents the difference between a transcript’s logFC and the overall logFC of the gene. For *Foxp1*, our results suggest a switch in usage of transcripts *Foxp1-216* and *Foxp1-225* between basal and LP cells. Previous work has reported transcript *Foxp1-225* to be exclusively expressed in basal cells and transcript *Foxp1-220* to be expressed at similar levels in both cell populations [7]. Our analysis additionally revealed that transcripts *Foxp1-214* and *Foxp1-216* are relatively higher expressed in LP cells compared to basal cells, and that transcript *Foxp1-210* is relatively higher expressed in basal cells compared to LP cells.

Among the 1,256 differentially used genes, we identified 656 genes with exactly two significantly differentially used transcripts exhibiting relative logFC of opposite signs. This finding suggests that switching expression among spliced transcripts is a prevalent phenomenon in the development of the mammary stem cell lineage. Notably, several genes identified by Milevskiy *et al*. [7] as having differentially expressed alternative start sites among cell populations were also statistically significant in our DTU analysis. One compelling example is *Eya2* (FDR = 1.4 × 10^−13^), the first two transcripts of which have different start sites and show mutually exclusive expression. Transcript *Eya2-202* (relative logFC = − 5.06, FDR = 2.0 × 10^−15^) is highly expressed in basal cells but completely absent in LP and ML. Transcript *Eya2-201* (relative logFC = 5.12, FDR = 4.4 × 10^−16^), on the other hand, is completely absent in basal but becomes expressed in LP and ML (Figure 5d).

The mammary data was also analyzed using *limma*, leading to almost the same number of significant results (1,260 DTU genes and 2,489 differentially used transcripts) and to the same top-ranked genes as discussed above.

## Discussion

In this article, we present fast and powerful pipelines for differential usage analysis of RNA-seq data with *limma* and *edgeR*. Our analysis pipeline employs the divided-count approach to *remove* the RTA overdispersion effect resulting from the transcript quantification step. An extensive simulation study demonstrates that both *limma* and *edgeR* pipelines, followed by differential usage testing via the *diffSplice* function, provides uniformly more powerful analyses than specialized DTU methods.

The *limma* pipeline makes use of the *voom* method, which transforms transcript divided-count data into log2-counts per million, estimates *voom* precision weights, and fits *limma* linear models while accounting for the loss of residual df caused by exact zeros, which is a common feature of RNA-seq data summarized at the transcript level. The *edgeR* pipeline makes use of the latest edgeR v4 quasi-negative-binomial methods with bias-corrected deviances for improved estimation of quasi-dispersions for lowly expressed transcripts [19]. We show that both *limma* and *edgeR* pipelines, when used with *diffSplice* for differential usage testing, control the false discovery rate (FDR) across all evaluated scenarios, including those with small numbers of replicates, at both the gene and transcript levels. We further demonstrate that the *diffSplice* pipelines are the only benchmarked methods to produce p-value distributions that are approximately uniform in null simulations. These pipelines leverage the fast and flexible statistical methods available in the *limma* and *edgeR* Bioconductor packages, placing our methods as a much faster and more powerful alternative than current DTU tools. Our case study of basal and LP cells from the steady-state mouse mammary gland identified several differentially used genes and transcripts that have previously been shown to be fundamental to the healthy development of the mammary gland stem cell lineage.

Our simulations show that divided counts, previously proposed for DTE [5], are equally effective for DTU. Dividing out the RTA dispersion tends to equalize the variability of different transcripts for the same gene and generally improves the fidelity of the divided counts to the familiar quadratic mean-variance relationship that is characteristic of bulk RNA-seq data [40, 19]. Dividing the counts also improved the performance of *DEXSeq* which, like *edgeR*, is based on negative binomial modelling of the transcript counts.

Our study focused on analysis of transcript counts, but we also made comparison with *BANDITS*, which infers DTU directly from the equivalence class counts and therefore can incorporate the uncertainty arising from RTA into the likelihood function. In principle, the equivalence class counts should contain complete information, but fully leveraging this information is computationally and statistically challenging. Our simulations show that the *diffSplice* pipelines are not only much simpler and faster but also achieved better statistical power and, mostly importantly, controlled the FDR far more accurately. *BANDITS* relies on *DRIM-Seq* to estimate genewise precision parameters, so it is still dependent on the transcript counts as well as on the equivalence class counts. *BANDITS* is also limited to two-group comparisons or oneway tests. It cannot accommodate covariates, or additive factors, or general treatment contrasts.

Overall, our simulations highlight the *edgeR* DTU pipeline combined with gene-level Simes p-values as the most powerful approach for detecting transcriptional differential usage in RNA-seq data. However, *limma* was a close competitor and, in more complex experimental settings, the added flexibility of the *limma* linear modeling framework may offer important advantages. Such scenarios include, for example, experimental designs with blocking factors where sampling units are correlated with an associated effect that must be accounted for as random effects, or when sample-specific quality weights must be estimated and accounted for during the differential analysis. Finally, our simulation results show that both *limma* and *edgeR* pipelines are robust to independent filtering, with efficient DTU analyses that are powerful to detect differential usage even from very lowly expressed transcripts.

The simulations presented in this article were designed to emulate small-scale RNA-seq experiments from model organisms with low levels of biological variability, such as those derived from genetically identical mice or cell lines. However, large-scale RNA-seq experiments with substantial sample sizes from human studies are also common. Our previous work has demonstrated the excellent performance of the divided-count approach for differential transcript expression in the context of larger samples sizes and larger levels of biological variability [5, 6], which suggests that the *diffSplice* pipelines will perform effectively in such scenarios as well, with the added advantage of computational scalability over current methods, owing to the efficient implementation of statistical methods in *limma* and *edgeR*.

Our simulations included two DTU scenarios whereby either one or two transcripts were differentially expressed for a given gene. This allowed us to show that the *diffSplice* Simes test is more powerful than the *diffSplice* F-test in the first scenario where just one transcript has a different pattern to the other transcripts for the same gene, whereas the *diffSplice* F-test is more powerful than the Simes test in the second scenario with two transcripts switching in a balanced way.

Our simulations make essentially no distributional assumptions except that the true TPM of each transcript vary between biological replicates with a constant coefficient of variation. The true TPM values were assumed to follow gamma distributions, but other distributions with constant coefficient of variation, such as log-normal, give similar results [5].

The simulations generated a random biological CV independently for each transcript, even though empirical evidence suggests that transcripts for the same gene tend to have similar CVs. The *limma* and *edgeR diffSplice* methods assume consistent dispersion for all transcripts of the same gene, but still performed strongly in the simulations, showing that they are robust to unequal dispersions. Nevertheless, it is reasonable to expect that a more elaborate and realistic simulation that allowed for BCV consistency within genes would improve the performance of *limma* and *edgeR* even more relative to other methods.

Another complication not included the simulation is the possibility of PCR duplicates in the RNA sequencing. In the *edgeR* context, PCR duplication tends to increase the quasi-dispersions of the affected transcripts [19]. The *limma* and *edgeR diffSplice* methods both allow for technical as well as biological variation, so both the methods should adapt smoothly to that complication, if present.

Another limitation of the current study is that we have not explored the effects of unannotated transcripts on any of the pipelines, and this remains an issue to be explored in the future.

Altogether, results from our simulation study demonstrate that *limma* and *edgeR* outperformed other specialized and popular DTU methods. Specifically, *limma* and *edgeR* were the only methods to correctly control the FDR in all scenarios while detecting more truly DTU genes and transcripts than other tools. Our simulation results indicate that the *edgeR* DTU pipeline with gene-level Simes p-values rank as the most powerful method to assess transcriptional differential usage from RNA-seq data. Yet, DTU analyses of complex real datasets may benefit from the extra flexibility of the *limma* linear modelling framework, such as in scenarios where sampling units are not independent and must be accounted for as random effects or when sample-specific quality weights must be estimated.

## Acknowledgements

We wish to acknowledge other team members who have worked on the related problem of differential exon usage over the past 15 years, including Davis McCarthy, who wrote *edgeR*’s *spliceVariants* function, and Charity Law, who wrote an earlier version of *limma*’s *diffSplice* function.

## Author contributions

P.B. designed the simulations, undertook the analyses, and wrote the paper. L.C. developed the *diffSplice* method for *edgeR* and the associated quasi-likelihood methods in *edgeR* v4, and coauthored the *fitFDistUnequalDF1* function in *limma*. M.L. developed the *voomLmFit* method. Y.C. developed and maintained *edgeR* software, including the earlier *diffSpliceDGE* method, originating some novel ideas that are preserved in the current function. G.S. conceived and supervised the project, co-authored and maintained software, and finalized the manuscript. All authors contributed to and checked the final manuscript.

## Data availability

The *limma, edgeR* and *Rsubread* packages are freely available from https://bioconductor.org/packages/limma, https://bioconductor.org/packages/edgeR, and https://bioconductor.org/packages/Rsubread respectively. Data and code to reproduce the results presented in this article are available from https://github.com/plbaldoni/DTU-code. Software versions used in the paper were *BANDITS*: 1.20.0, *DEXSeq*: 1.50.0, *DRIMSeq*: 1.32.0, *edgeR*: 4.5.9, *limma*: 3.63.9, *R*: 4.4.1, *Rsubread* : 2.18.0, *Salmon*: 1.10.2, *satuRn*: 1.12.0.

The example RNA-seq data analyzed in this article was downloaded from https://www.ncbi.nlm.nih.gov/geo/query/acc.cgi?acc=GSE227748.

## Supplementary data

Supplementary figures and tables are available in the file supp.pdf.

