## Supplementary Material for "Dividing out quantification uncertainty enables assessment of differential transcript usage with limma and edgeR"

### Supplementary materials for
### "Dividing out quantification uncertainty enables fast and accurate assessment of differential transcript usage with limma and edgeR"

Pedro L. Baldoni[1,2,†], Lizhong Chen[1,2,†], Mengbo Li[1,2],
Yunshun Chen[1,2,3], and Gordon K. Smyth[1,2,*]

28 September 2025

#### Contents

**1 Supplementary Figures** **2**

**2 Supplementary Tables** **19**

[1]Bioinformatics and Computational Biology Division, WEHI, Parkville, VIC 3052, Australia,
[2]Department of Medical Biology, The University of Melbourne, Parkville, VIC 3010, Australia,
[3]ACRF Cancer Biology and Stem Cells Division, WEHI, Parkville, VIC 3052, Australia,
[†]These authors contributed equally to this work.

### 1 Supplementary Figures

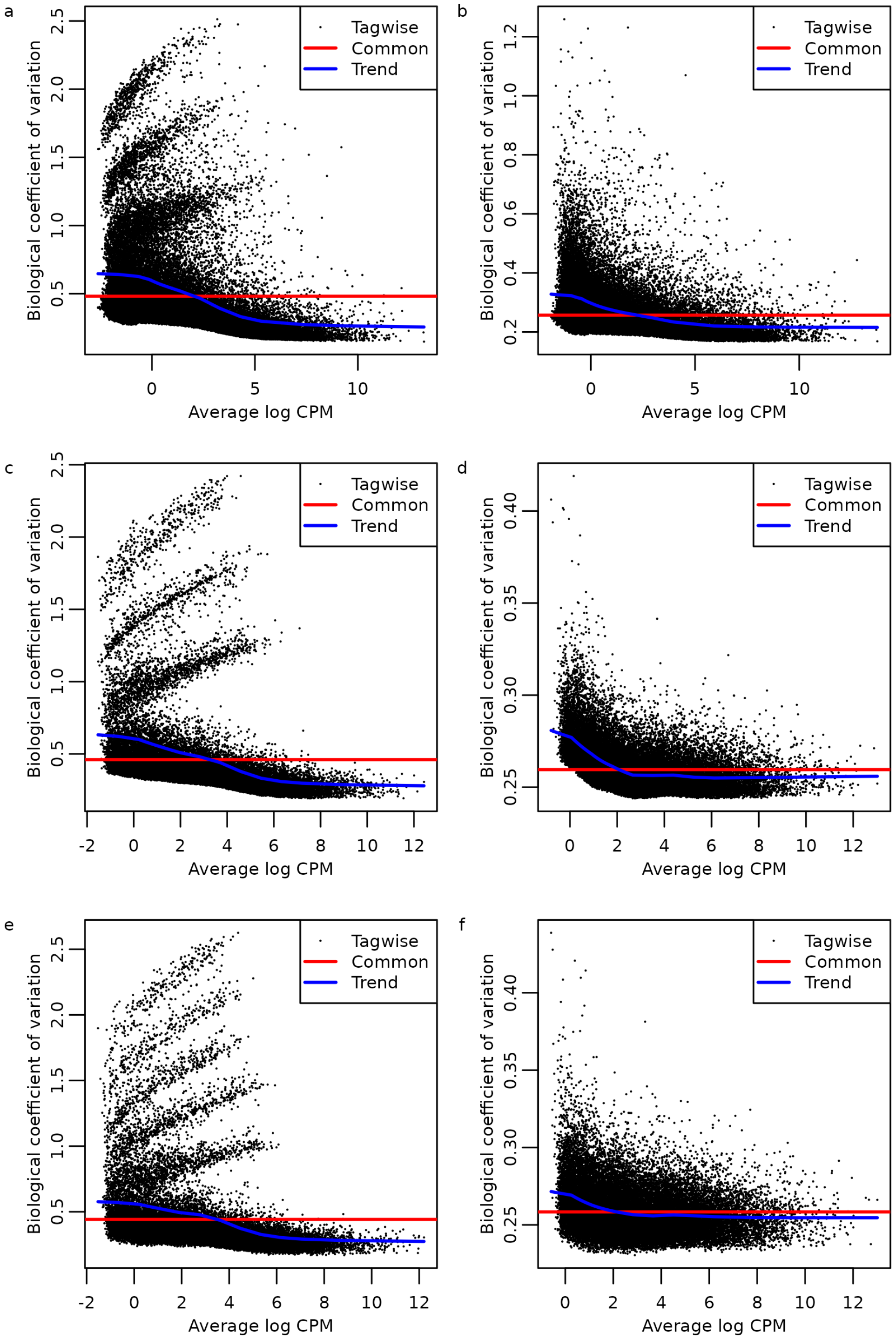


Figure S1: BCV plots of transcript-level counts from real and simulated data. Panels (a) and (b) show results from RNA-seq libraries of basal and luminal progenitor cells from the the mouse mammary gland data before and after count scaling, respectively. Panels (c) and (d) show results from a representative simulated mouse RNA-seq experiment with $n = 3$ samples per group before and after count scaling, respectively. Panels (e) and (f) show results from a representative simulated mouse RNA-seq experiment with $n = 5$ samples per group before and after count scaling, respectively. Standard expression filtering by *filterByExpr* but no annotation filtering.

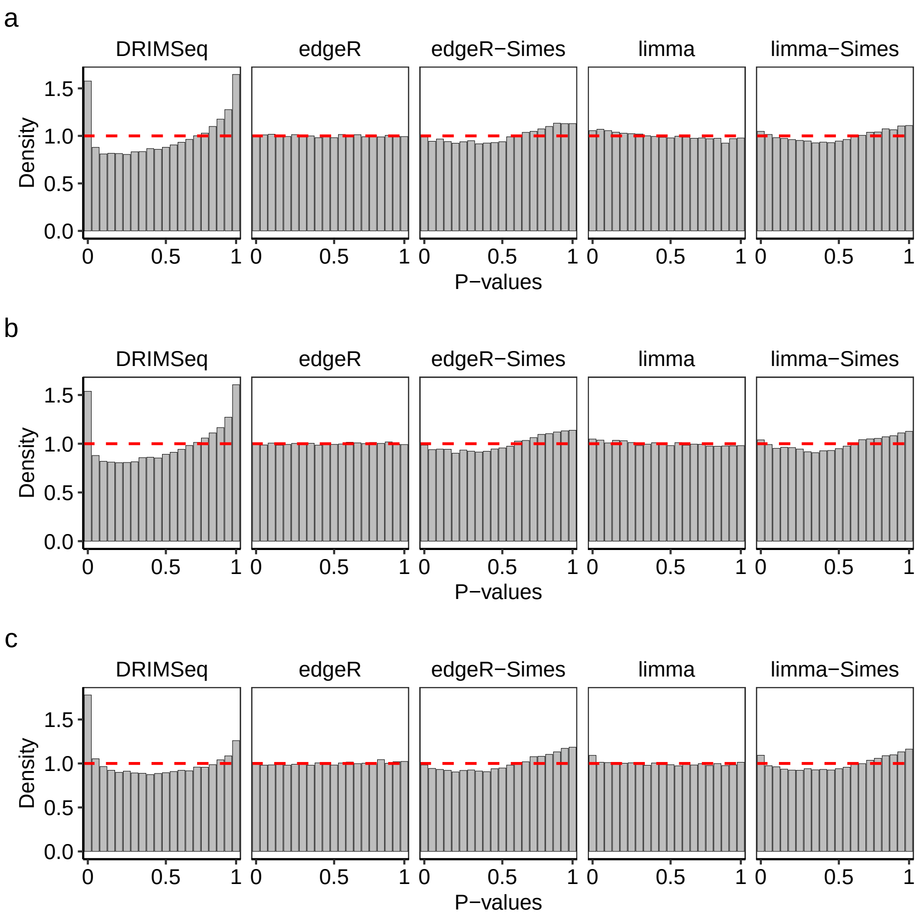


Figure S2: Density histograms of gene-level p-values in the null simulation. Dashed line indicates the expected uniform distribution. Panels (a)–(c) show results with three, five, and ten samples per group. Methods *DEXSeq* and *satuRn* do not report gene-level p-values and, therefore, are omitted in this figure. Raw counts were used by *DEXSeq*, *DRIMSeq*, and *satuRn*. Results are averaged over 20 simulated datasets.

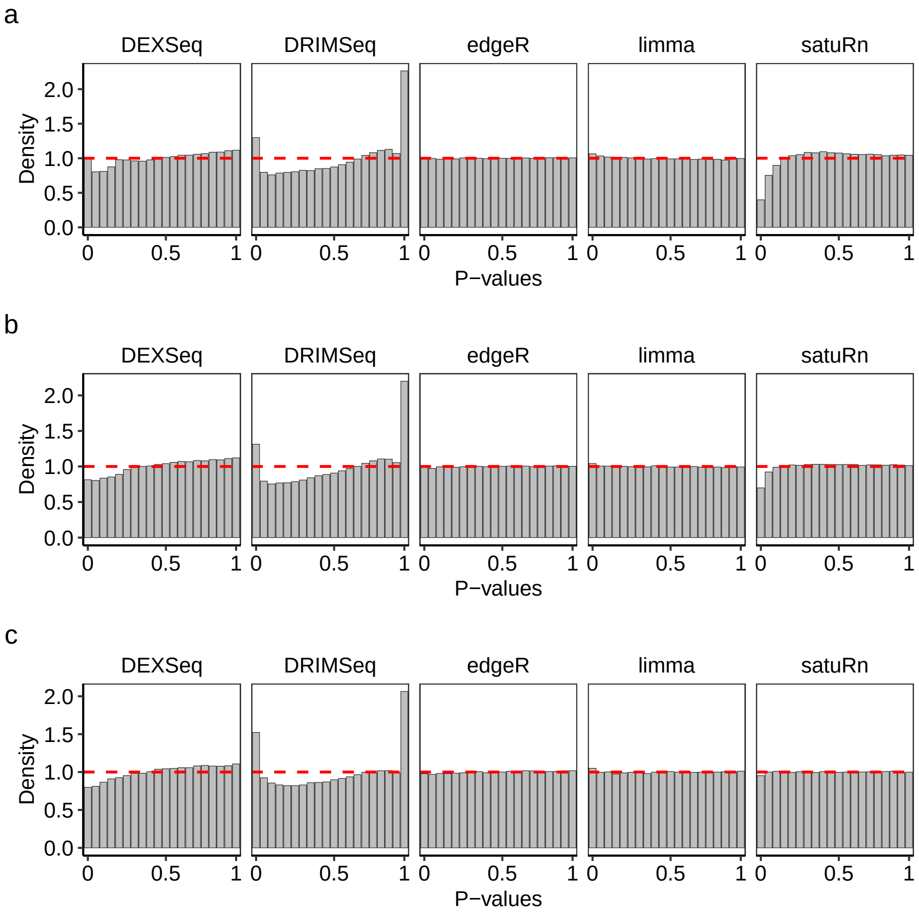


Figure S3: Density histograms of transcript-level p-values in the null simulation. Dashed line indicates the expected uniform distribution. Panels (a)–(c) show results with three, five, and ten samples per group. Raw counts were used by *DEXSeq*, *DRIMSeq*, and *satuRn*. Results are averaged over 20 simulations.

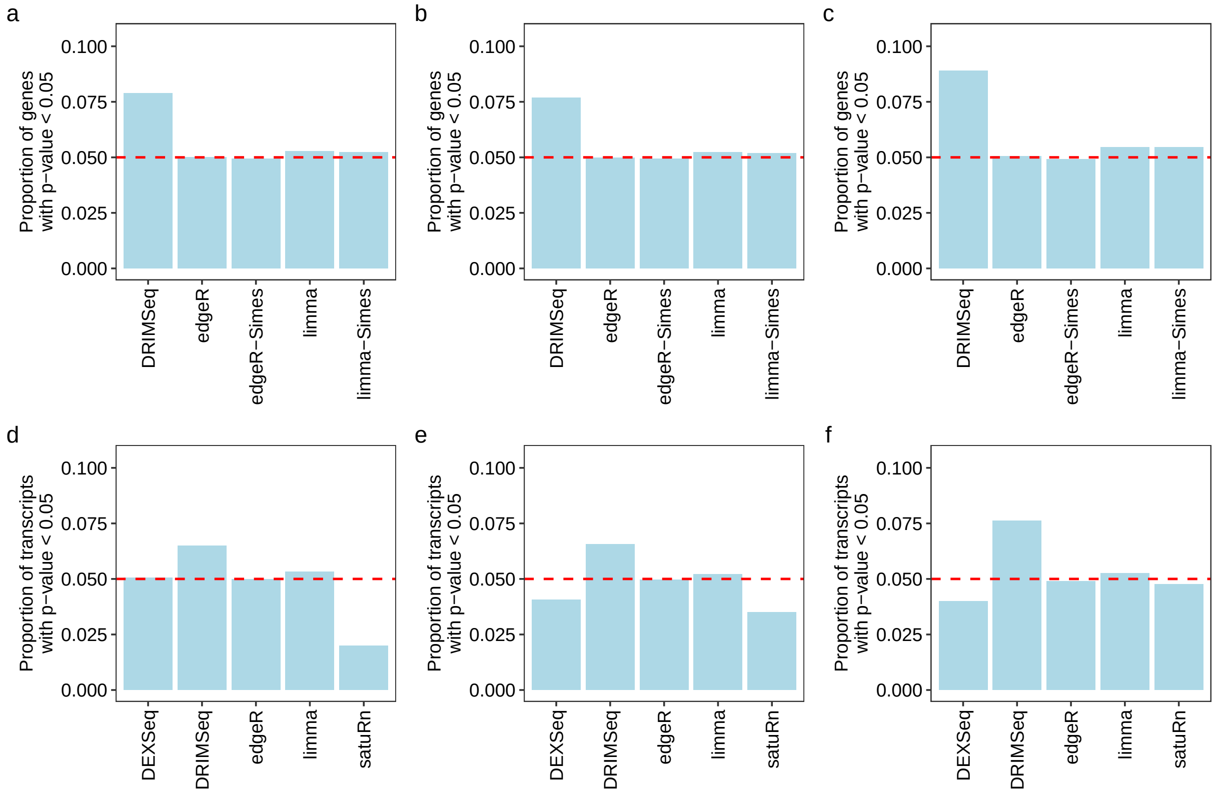


Figure S4: Type 1 error rate at 5% level. Proportion of genes or transcripts with unadjusted p-values $< 0.05$ in the null simulation. The dashed line indicates the expected proportion of 0.05. Panels (a)–(c) show results at the gene-level. Panels (d)–(f) show results at the transcript-level. In (a) and (d), scenario with three samples per group. In (b) and (e), scenario with five samples per group. In (c) and (f), scenario with ten samples per group. Methods *DEXSeq* and *satuRn* do not report gene-level raw p-values and, therefore, are omitted from panels (a)–(c). Raw counts were used by *DEXSeq*, *DRIMSeq*, and *satuRn*. Results are averaged over 20 simulations.

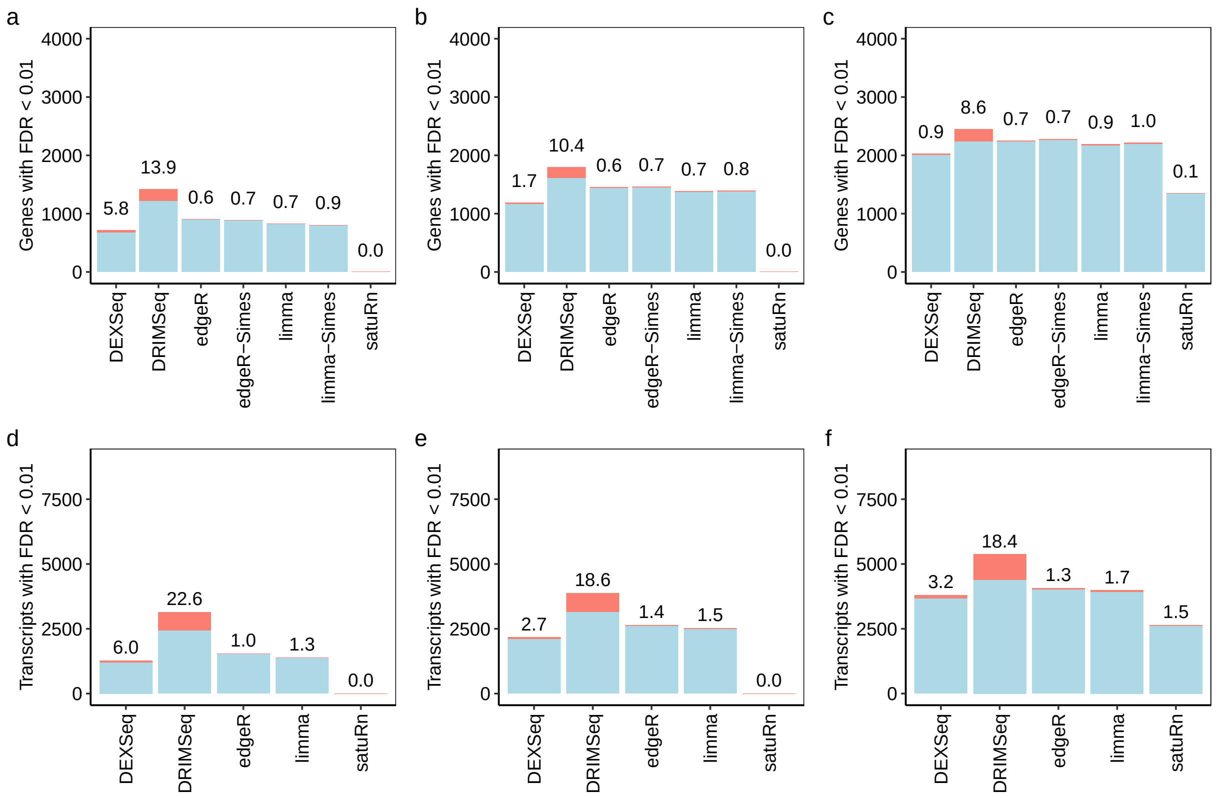


Figure S5: Numbers of true and false discoveries at 1% FDR. Stacked barplots showing the number of true (blue) and false (red) positive differentially used genes and transcripts at nominal 1% FDR. The observed FDR is shown as a percentage over each bar. Panels (a)–(c) show results at the gene-level with three, five, and ten samples per group, respectively. Panels (d)–(f) show results at the transcript-level with three, five, and ten samples per group, respectively. The y-axis scales match those of Figure 3 to enable easy comparison. *satuRn* consistently failed to detect differentially used genes and transcripts in scenarios with three samples per group. Raw counts were used by *DEXSeq*, *DRIMSeq*, and *satuRn*. Results are averaged over 20 simulated datasets.

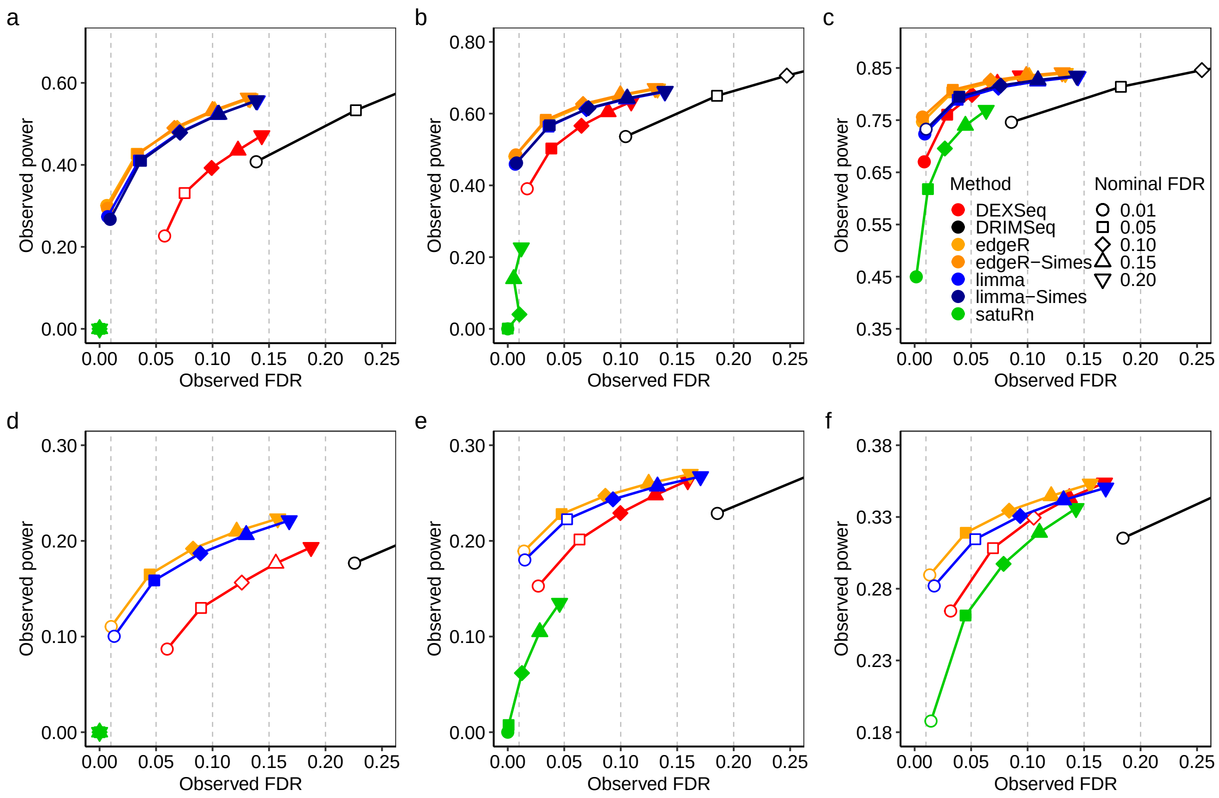


Figure S6: Power vs FDR. Observed power (true positive rate) over observed FDR (false discovery rate) for different methods and different nominal FDR threshold levels at the gene and transcript-levels. Colored lines connect data points from each method. Different point shapes represent different nominal FDR thresholds, and are filled if the observed FDR is less than the nominal value. Nominal FDR levels are indicated by dashed vertical lines. Panels (a)–(c) show results at the gene-level with three, five, and ten samples per group, respectively. Panels (d)–(f) show results at the transcript-level with three, five, and ten samples per group, respectively. The method *satuRn* consistently failed to detect differentially used genes and transcripts in scenarios with three samples per group, and under a nominal 0.01 FDR threshold with five samples per group. Raw counts were used by *DEXSeq*, *DRIMSeq*, and *satuRn*. Results are averaged over 20 simulations.

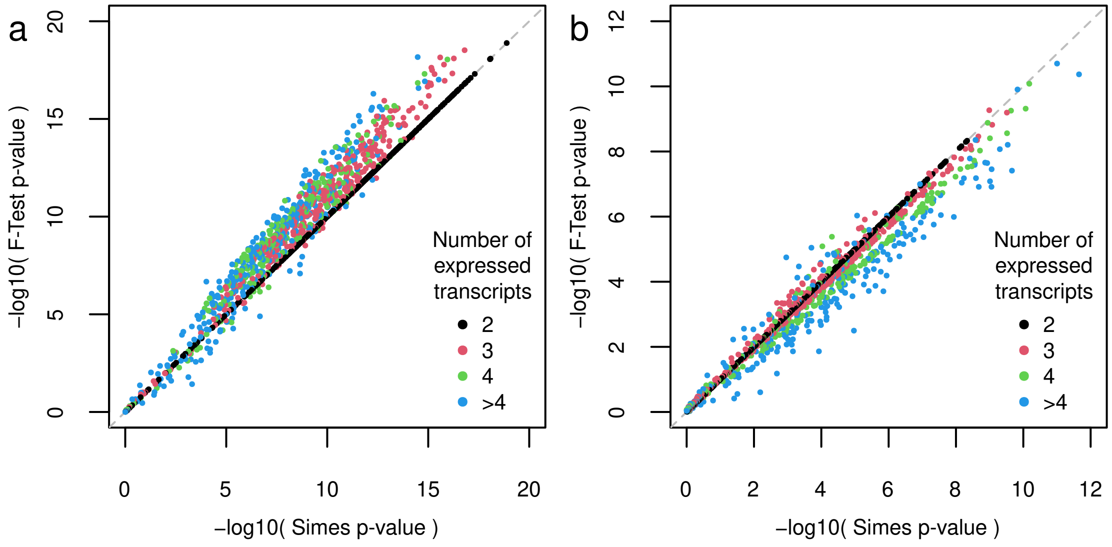


Figure S7: F-test vs Simes p-values from *edgeR* for a representative simulated dataset with $n = 10$ samples per group. Panel (a) plots p-values for genes with two adjusted transcripts (the DTU-only genes) while panel (b) plots p-values for genes with a single adjusted transcript (the DGE/DTU genes). All points correspond to genes with true DTU, so that smaller p-values are better. The total number of transcripts per gene is indicated by colour. P-values are represented on the minus log10 scale.

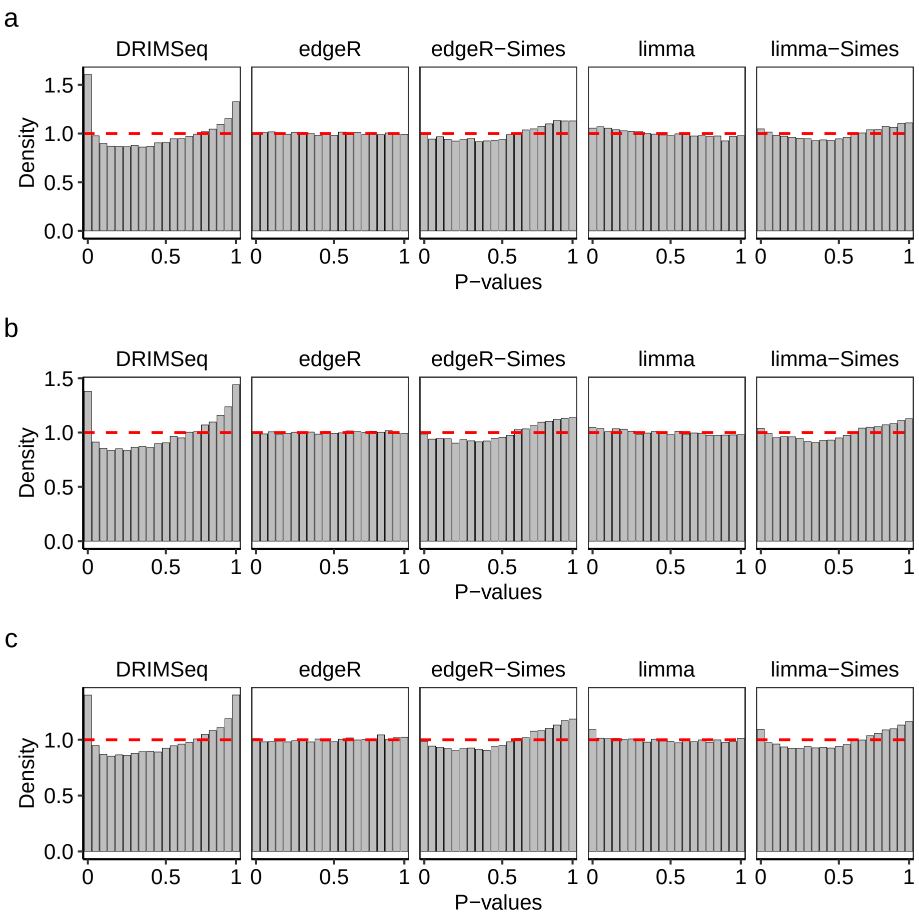


Figure S8: Density histograms of gene-level p-values in the null simulation. Dashed line indicates the expected uniform distribution. Panels (a)–(c) show results with three, five, and ten samples per group. Methods *DEXSeq* and *satuRn* do not report gene-level p-values and, therefore, are omitted in this figure. Divided counts were used by all methods. Results are averaged over 20 simulated datasets.

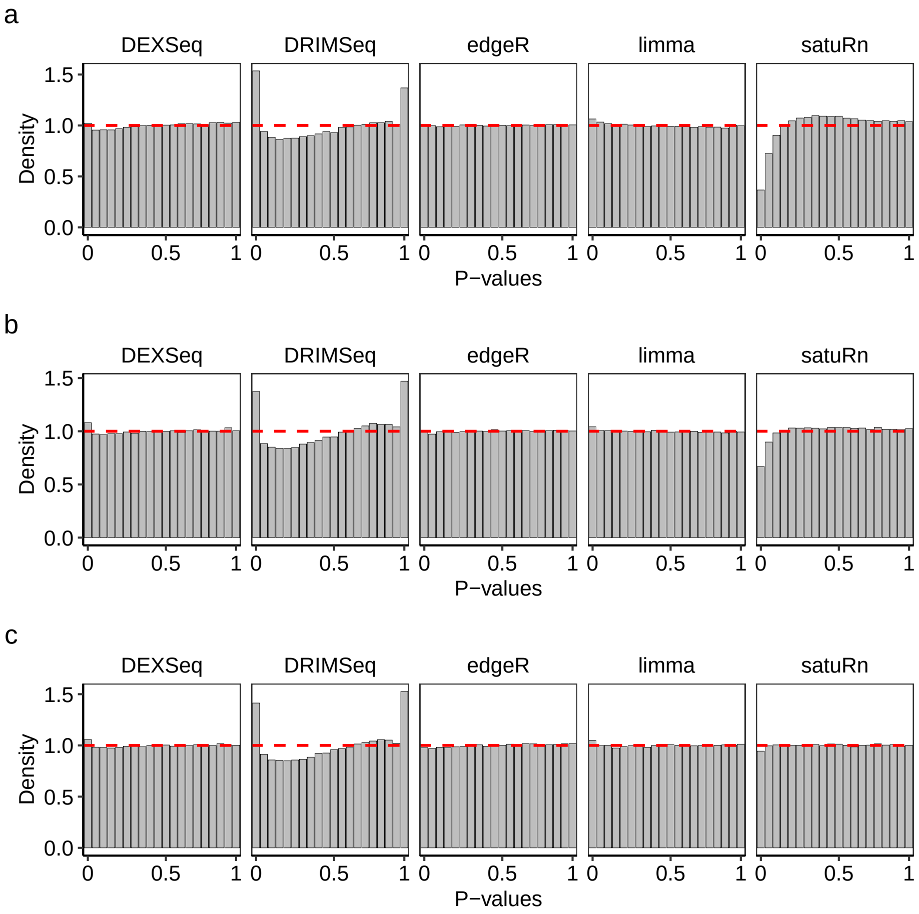


Figure S9: Density histograms of transcript-level p-values in the null simulation. Dashed line indicates the expected uniform distribution. Panels (a)–(c) show results with three, five, and ten samples per group. Divided counts were used by all methods. Results are averaged over 20 simulations.

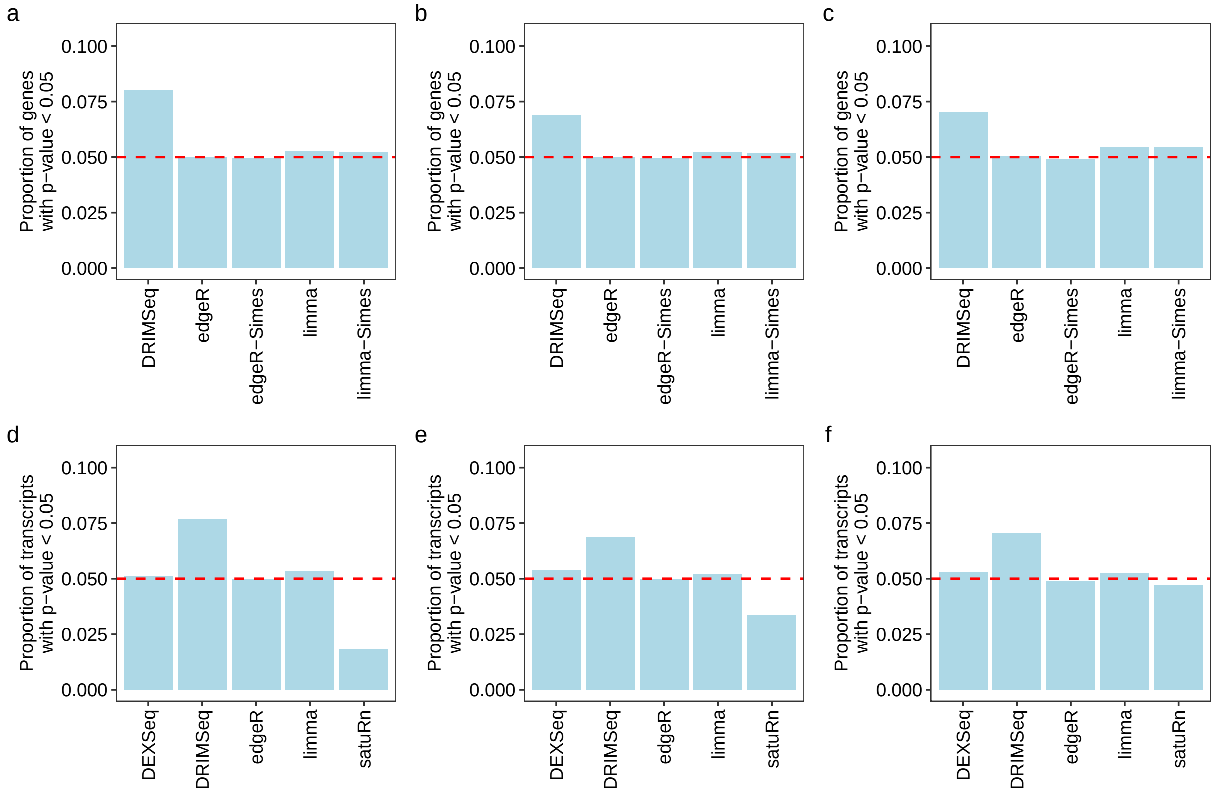


Figure S10: Type 1 error rate at 5% level. Proportion of genes or transcripts with unadjusted p-values $< 0.05$ in the null simulation. The dashed line indicates the expected proportion of 0.05. Panels (a)–(c) show results at the gene-level. Panels (d)–(f) show results at the transcript-level. In (a) and (d), scenario with three samples per group. In (b) and (e), scenario with five samples per group. In (c) and (f), scenario with ten samples per group. Methods *DEXSeq* and *satuRn* do not report gene-level raw p-values and, therefore, are omitted from panels (a)–(c). Divided counts were used by all methods. Results are averaged over 20 simulations.

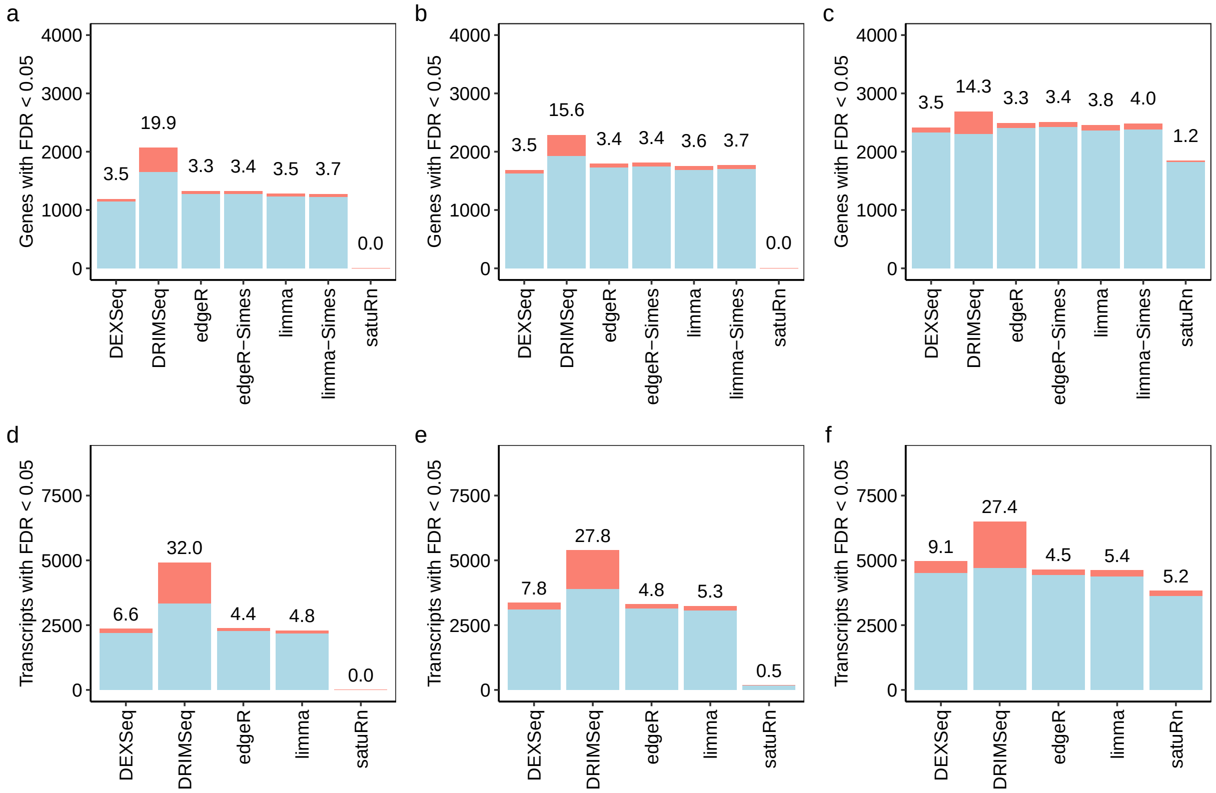


Figure S11: Numbers of true and false discoveries at 5% FDR. Stacked barplots show the number of true (blue) and false (red) positive differentially used genes and transcripts at nominal 5% FDR. The observed FDR is shown as a percentage over each bar. Panels (a)–(c) show results at the gene-level with three, five, and ten samples per group, respectively. Panels (d)–(f) show results at the transcript-level with three, five, and ten samples per group, respectively. The y-axis scales match those of Figures 3 and S5 to enable easy comparison. *satuRn* failed to detect differentially used genes or transcripts with three samples per group. Divided counts were used by all methods. Results are averaged over 20 simulations.

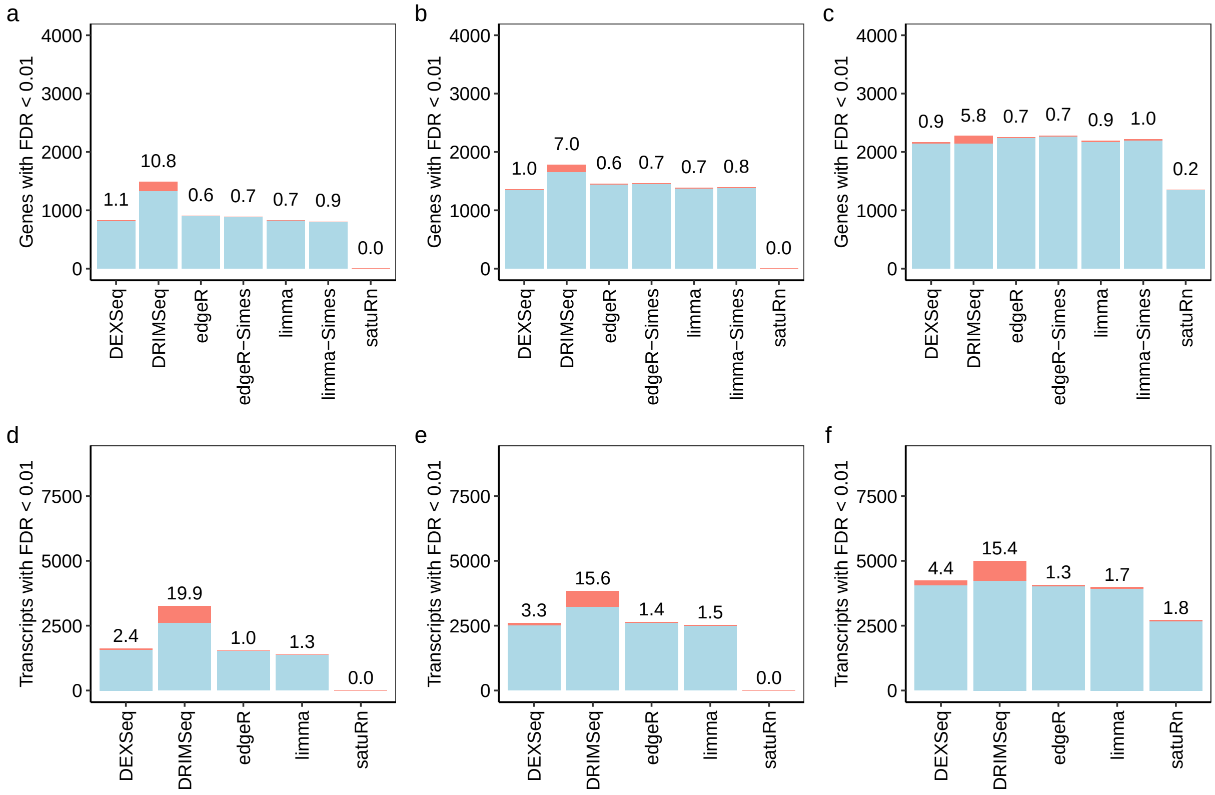


Figure S12: Numbers of true and false discoveries at 1% FDR. Stacked barplots showing the number of true (blue) and false (red) positive differentially used genes and transcripts at nominal 1% FDR. The observed FDR is shown as a percentage over each bar. Panels (a)–(c) show results at the gene-level with three, five, and ten samples per group, respectively. Panels (d)–(f) show results at the transcript-level with three, five, and ten samples per group, respectively. The y-axis scales match those of Figures 3, S5 and S11 to enable easy comparison. *satuRn* consistently failed to detect differentially used genes and transcripts in scenarios with three samples per group. Divided counts were used by all methods. Results are averaged over 20 simulated datasets.

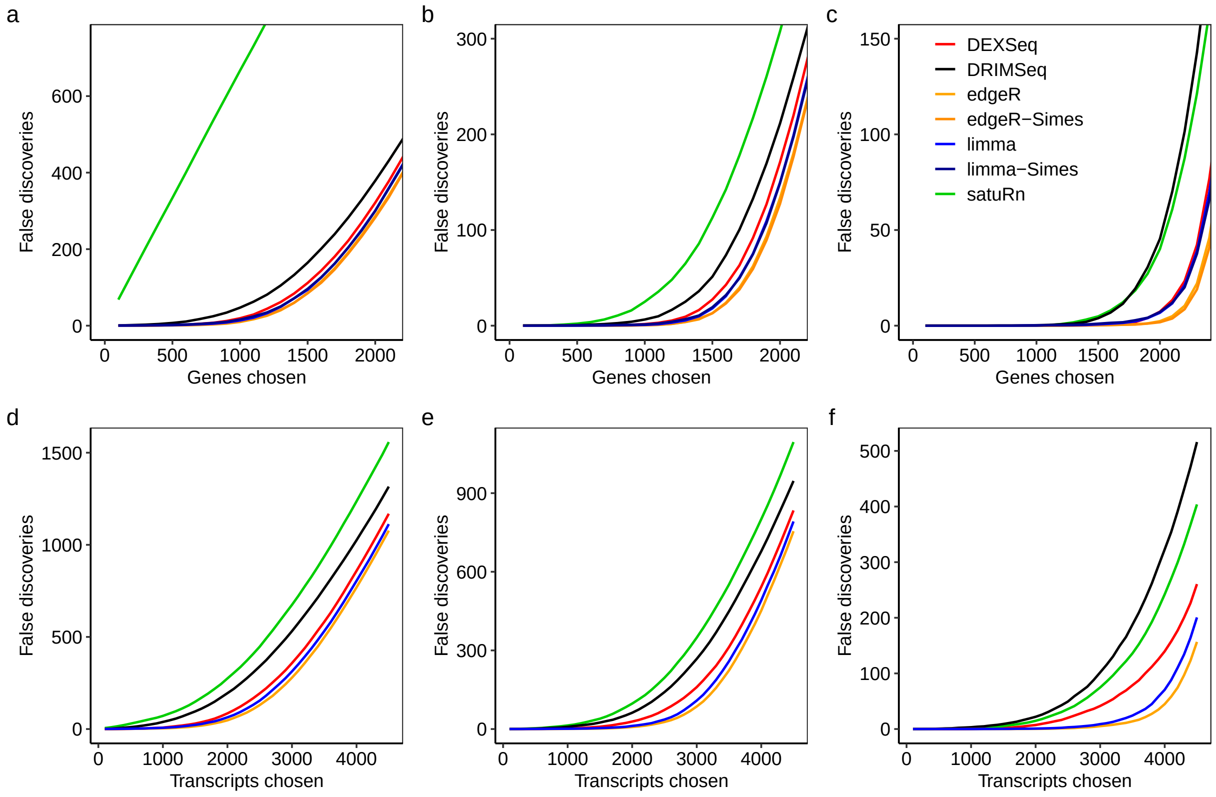


Figure S13: Cumulative false discoveries in ranked lists. The number of false discoveries is plotted for each method versus the number of genes or transcripts selected as differentially used. Panels (a)–(c) show gene-level results with three, five, and ten samples per group, respectively. Panels (d)–(f) show transcript-level results with three, five, and ten samples per group, respectively. For *DEXSeq* and *satuRn*, genes were ranked according to their adjusted p-values, as those methods do not provide unadjusted p-values by default. Divided counts were used by all methods. Results are averaged over 20 simulations.

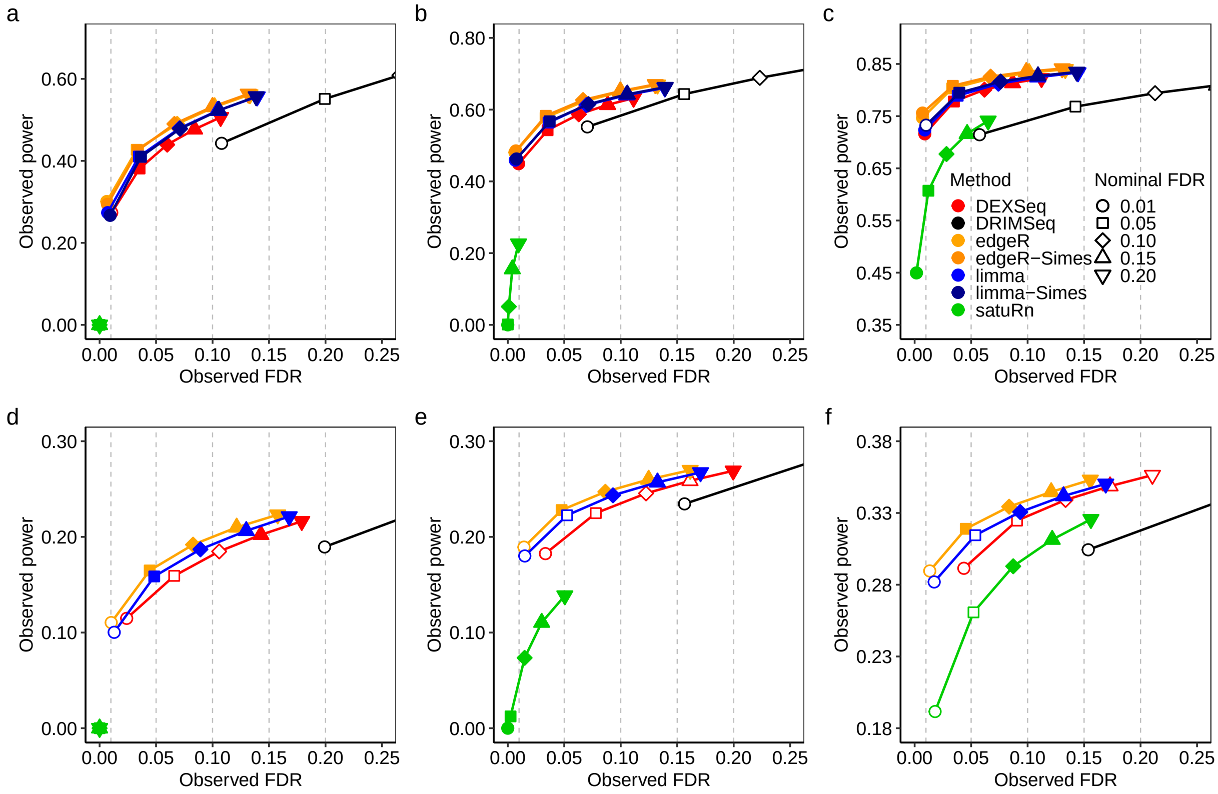


Figure S14: Power vs FDR. Observed power (true positive rate) over observed FDR (false discovery rate) for different methods and different nominal FDR threshold levels at the gene and transcript-levels. Colored lines connect data points from each method. Different point shapes represent different nominal FDR thresholds, and are filled if the observed FDR is less than the nominal value. Nominal FDR levels are indicated by dashed vertical lines. Panels (a)–(c) show results at the gene-level with three, five, and ten samples per group, respectively. Panels (d)–(f) show results at the transcript-level with three, five, and ten samples per group, respectively. The method *satuRn* consistently failed to detect differentially used genes and transcripts in scenarios with three samples per group, and under a nominal 0.01 FDR threshold with five samples per group. Divided counts were used by all methods. Results are averaged over 20 simulations.

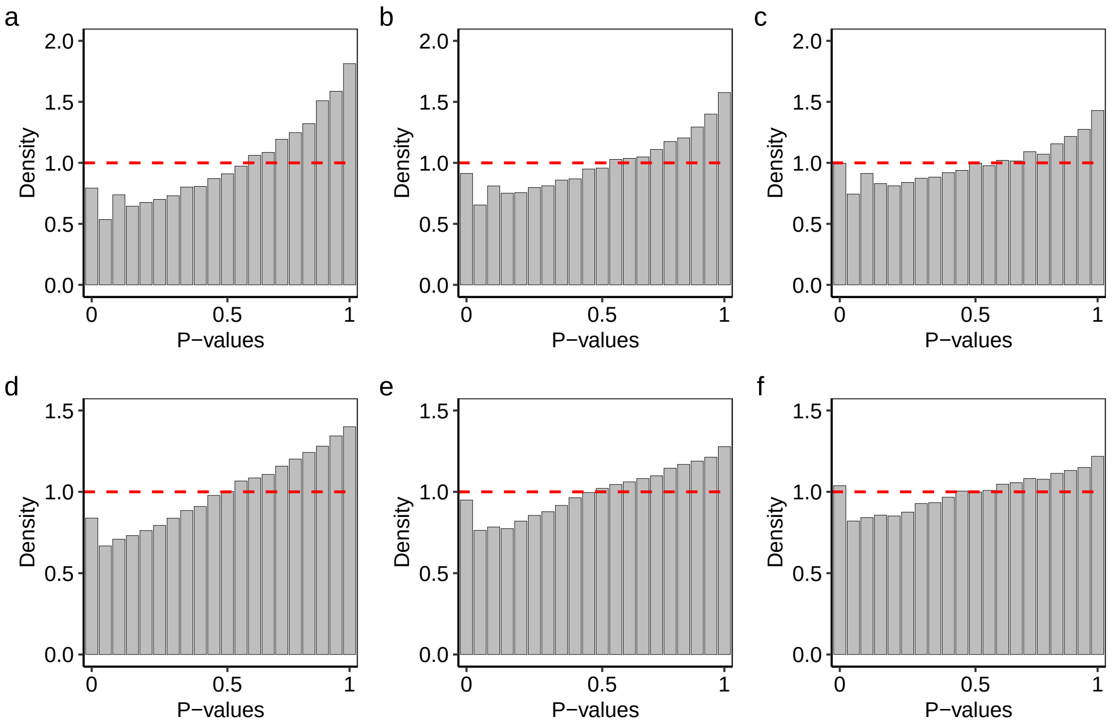


Figure S15: Density histograms of p-values from *BANDITS* for the null simulations. Dashed line indicates the expected uniform distribution. Panels (a)–(c) show gene-level results with three, five, and ten samples per group. Panels (d)–(f) show transcript-level results with three, five, and ten samples per group. Results are averaged over 5 simulated datasets.

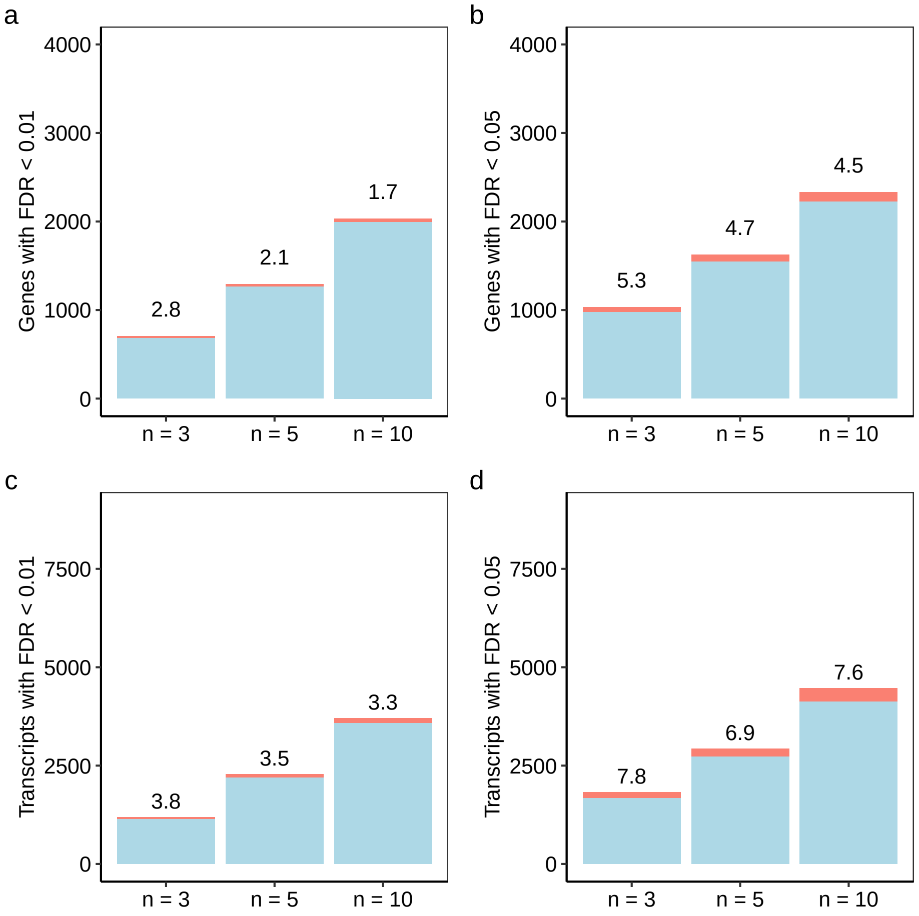


Figure S16: Numbers of true and false discoveries from *BANDITS*. Stacked barplots showing the number of true (blue) and false (red) positive differentially used genes and transcripts for scenarios with $n = 3$, $n = 5$, and $n = 10$ samples per group. The observed FDR is shown as a percentage over each bar. Panels (a)–(b) show results at the gene-level with 0.01 and 0.05 FDR cutoff, respectively. Panels (c)–(d) show results at the transcript-level with 0.01 and 0.05 FDR cutoff, respectively. The y-axis scales match those of Figures 3, S5, S11 and S12 to enable easy comparison. Results are averaged over 5 simulated datasets.

### 2 Supplementary Tables

Table S1: Strict vs lenient filtering at gene level. Number of true and false discoveries at gene-level with 0.05 FDR cutoff for *edgeR* and *limma* with strict and lenient filtering. The strict criterion represents the *filterByExpr* function with default values *min.count = 10* and *min.total.count = 15*. The lenient criterion represents *filterByExpr* with customized values *min.count = 1* and *min.total.count = 5*. Results are averaged over 20 simulations.

| | | Strict | | | Lenient | | |
|---|---|---|---|---|---|---|---|
| Samples per group | Method | TP | FP | FDR | TP | FP | FDR |
| Three | edgeR | 1281.00 | 43.55 | 3.3 | 1255.05 | 55.05 | 4.2 |
| | edgeR-Simes | 1277.45 | 44.95 | 3.4 | 1258.75 | 56.35 | 4.3 |
| | limma | 1231.45 | 44.50 | 3.5 | 1280.60 | 70.60 | 5.2 |
| | limma-Simes | 1229.55 | 46.75 | 3.7 | 1288.30 | 69.90 | 5.1 |
| Five | edgeR | 1733.55 | 60.10 | 3.4 | 1922.40 | 74.95 | 3.8 |
| | edgeR-Simes | 1748.40 | 60.95 | 3.4 | 1944.30 | 75.10 | 3.7 |
| | limma | 1691.65 | 63.50 | 3.6 | 1874.20 | 88.90 | 4.5 |
| | limma-Simes | 1702.55 | 65.75 | 3.7 | 1889.50 | 92.25 | 4.7 |
| Ten | edgeR | 2409.80 | 82.90 | 3.3 | 2561.35 | 104.80 | 3.9 |
| | edgeR-Simes | 2424.40 | 84.85 | 3.4 | 2582.50 | 103.85 | 3.9 |
| | limma | 2367.05 | 93.80 | 3.8 | 2490.70 | 117.95 | 4.5 |
| | limma-Simes | 2384.60 | 98.30 | 4.0 | 2512.00 | 123.55 | 4.7 |

*Note:*

TP: true positive genes. FP: false positive genes. FDR: false discovery rate.

Table S2: Strict vs lenient filtering at transcript level. Number of true and false discoveries at transcript-level with 0.05 FDR cutoff for *edgeR* and *limma* with strict and lenient filtering. The strict criterion represents the *filterByExpr* function with default values *min.count = 10* and *min.total.count = 15*. The lenient criterion represents *filterByExpr* with customized values *min.count = 1* and *min.total.count = 5*. Results are averaged over 20 simulations.

| | | Strict | | | Lenient | | |
|---|---|---|---|---|---|---|---|
| Samples per group | Method | TP | FP | FDR | TP | FP | FDR |
| Three | edgeR | 2274.15 | 105.55 | 4.4 | 2176.45 | 109.85 | 4.8 |
| | limma | 2188.30 | 111.30 | 4.8 | 2233.80 | 145.60 | 6.1 |
| Five | edgeR | 3149.60 | 157.80 | 4.8 | 3454.25 | 156.80 | 4.3 |
| | limma | 3074.35 | 170.40 | 5.3 | 3363.65 | 197.15 | 5.5 |
| Ten | edgeR | 4432.85 | 210.20 | 4.5 | 4675.50 | 224.00 | 4.6 |
| | limma | 4370.90 | 248.10 | 5.4 | 4564.35 | 272.25 | 5.6 |

*Note:*

TP: true positive transcripts. FP: false positive transcripts. FDR: false discovery rate.
